# Podoplanin shapes epidermal morphogenesis and coordinates stem cell and epidermal-dermal responses during inflammatory remodelling

**DOI:** 10.64898/2026.11.03.760123

**Authors:** Rocío Jarcovsky, Sara Temprano-Carazo, Ana Bravo, Santiago Garcés-Fernández, Blanca Gómez-Barahona, Marina Gilolmo, Silvia Roca, Diego Megías, Susana Barroso, Paloma Fernández, Virginia García-García, Javier Moratinos, Susana Álvarez, Lucía Montero-Montero, Esther Grueso, Jaime Millán, Steve P Watson, Ester Martín-Villar

**Affiliations:** Faculty of Experimental Sciences. Institute of Life Sciences. Universidad Francisco de Vitoria, Madrid, Spain; Department of Anatomy, Animal Production, and Veterinary Clinical Sciences. Laboratory of Pathology Phenotyping of Genetically Engineered Mice. Faculty of Veterinary Medicine. University of Santiago de Compostela, Lugo, Spain; Advanced Optical Microscopy Unit. Instituto de Salud Carlos III, Madrid, Spain; Nemesio Díez Institute of Applied Molecular Medicine. Universidad San Pablo-CEU, Madrid, Spain; Centro de Biología Molecular Severo Ochoa, CSIC-UAM, Madrid, Spain; Institute of Biomedical Research. Department of Cardiovascular Sciences. School of Medical Sciences. College of Medicine and Health. University of Birmingham, Birmingham, United Kingdom; Instituto de Investigación Sanitaria del Hospital Universitario La Paz (IdiPAZ), Madrid, Spain

## Abstract

Skin development and adaptation to inflammatory stress require keratinocytes to coordinate changes in cell behaviour with tissue organisation. Cell adhesion and the cortical cytoskeleton support the structural plasticity underlying these responses. Here, using complementary mouse models of global podoplanin deletion and conditional deletion in basal keratinocytes driven by keratin 5-Cre (K5Cre), we identify a role for the small transmembrane glycoprotein podoplanin in maintaining basal layer organisation and coordinating differentiation with barrier acquisition during development, as well as in inflammatory skin remodelling. Podoplanin expression was dynamically regulated during epidermal morphogenesis, becoming enriched in basal epithelial compartments before declining postnatally in the interfollicular epidermis. Both deletion models exhibited basal layer disorganisation, altered division orientation and increased epidermal thickness. Premature or expanded expression of differentiation markers was accompanied by delayed barrier acquisition, indicating that podoplanin loss uncouples differentiation marker expression from functional maturation. Barrier function subsequently recovered despite persistent histological abnormalities. In primary keratinocytes, podoplanin deficiency disrupted junctional actin organisation, reduced ERM phosphorylation and impaired junctional enrichment of pERM and doubly phosphorylated myosin light chain 2 (ppMLC2), despite largely preserved recruitment of adhesion proteins. In adult skin, K5Cre-mediated deletion altered differentiation marker dynamics following topical application of the proinflammatory agent 12-O-tetradecanoylphorbol-13-acetate (TPA), while the epidermal thickening response remained largely comparable to controls. These epidermal changes were accompanied by altered dermal thickness dynamics and an absence of TPA-induced podoplanin expression in dermal fibroblasts. Furthermore, TPA reduced the number of BrdU-retaining cells in the interfollicular epidermis of controls but not K5Cre-mediated conditional podoplanin knockout mice. This difference is consistent with impaired activation or mobilisation of slow-cycling cells, suggesting altered responsiveness within the epidermal stem and progenitor compartment. Together, these findings establish keratinocyte-expressed podoplanin as a regulator of epidermal organisation and functional maturation, with effects extending beyond the epithelium to shape the dermal response to inflammatory challenge.

## INTRODUCTION

The formation of a functional epidermis requires the coordinated proliferation, stratification and differentiation of basal keratinocytes. During development, these processes transform a simple epithelial layer into a multilayered barrier, while in adult skin they sustain tissue renewal and adaptation to injury or inflammation. Cell polarity, cytoskeletal organisation and cell-cell and cell-matrix adhesion contribute to this coordination by shaping basal cell architecture, division orientation and the transition towards suprabasal differentiation (Gadre et al., 2024; Miroshnikova et al., 2018; Simpson et al., 2011; Yin et al., 2023). These cellular processes help regulate the balance between renewal and differentiation of epidermal stem and progenitor populations in the interfollicular epidermis (IFE) and hair follicles (HFs), enabling tissue maintenance and adaptation to changing local demands (Gadre et al., 2024; Ishikawa et al., 2025). Dysregulation of epidermal stem cell function has been associated to a range of pathological conditions, including defective wound healing, chronic inflammatory skin diseases, and skin carcinogenesis (Gonzales and Fuchs, 2017; Hur, 2024). As an interface between keratinocytes and their local environment, cell-surface proteins offer a basis for understanding how neighbouring cells and the extracellular matrix influence epidermal organisation and differentiation.

Podoplanin (PDPN) is a small transmembrane glycoprotein involved in the regulation of cell motility, adhesion and epithelial plasticity (Quintanilla et al., 2019). Its expression is detected in several tissues, including the lymphatic endothelium, lung and heart, where it plays important roles during development. Accordingly, podoplanin-deficient mice display severe developmental abnormalities and die either during embryogenesis or shortly after birth due to defects affecting the cardiovascular and respiratory systems (Mahtab et al., 2008; Schacht et al., 2003). Mice overcoming these defects finally die within the first two postnatal weeks probably affected by the lymphatic system dysfunctions or other yet unknown causes. In the adult organism, podoplanin participates in diverse physiological processes such as lymphangiogenesis, platelet activation/aggregation, and innate immune responses (Astarita et al., 2012; Quintanilla et al., 2019). In addition, aberrant podoplanin expression has been consistently associated with tumour progression and chronic inflammatory diseases, including those affecting the skin (Asai et al., 2022) and has also been involved in tissue remodelling processes such as those occurring after cardiac injury (Cimini and Kishore, 2021) .

In adult skin, podoplanin is expressed by lymphatic endothelial cells (LECs) and selected epithelial compartments, including the basal layer of sebaceous glands and the outer root sheath (ORS) of growing hair follicles, but is barely detectable in the interfollicular epidermis (IFE) under homeostatic conditions (Gandarillas et al., 1997; Honma et al., 2012; Yoon et al., 2019). Its expression is induced in basal interfollicular keratinocytes and dermal fibroblasts during inflammation, tissue repair and carcinogenesis, and has been associated with the progression of cutaneous and head and neck squamous cell carcinomas (SCCs) (Gandarillas et al., 1997; Honma et al., 2012; Martín-Villar et al., 2005; Toll et al., 2012). In keratinocytes, this induction can be driven by TGF-β through Smad signalling and by IFN-γ, IL-6 and IL-22 through JAK-STAT pathways (Honma et al., 2012), while Fos/AP-1-dependent transcriptional regulation has been described during skin carcinogenesis (Durchdewald et al., 2008). Among its molecular functions, podoplanin regulates actin cytoskeleton organisation through its interaction with ezrin, radixin and moesin (ERM) and the modulation of Rho GTPase-family signalling. Studies in cultured epithelial and SCC cells have linked these activities to changes in cell shape, adhesion, motility and invasion (Martín-Villar et al., 2015, 2006). Its contribution to epidermal physiology *in vivo*, however, remains less clearly defined. Previous studies using epithelial deletion models have focused primarily on adult skin. K5Cre-mediated deletion altered hair follicle cycling but produced no overt abnormalities in the interfollicular epidermis under homeostatic conditions (Yoon et al., 2019), while no impairment of gross re-epithelialisation was detected following acute skin wounding in mice with K14Cre-mediated deletion (Baars et al., 2015).

These observations leave unresolved whether podoplanin contributes to epidermal architecture during development or subsequent remodelling. Its ability to regulate the cytoskeleton and cell adhesion provides a rationale for examining epidermal morphogenesis, when these processes must be coordinated with stratification, differentiation and barrier acquisition. Likewise, its induction during inflammation raises the question of whether the expression of podoplanin in keratinocytes contributes to the tissue response. Studying these contexts may therefore reveal functions that are less evident in adult epidermis under steady-state conditions, where podoplanin is barely detectable. In this study, we used complementary global and K5Cre-mediated conditional knockout mouse models to define how podoplanin contributes to epidermal organisation and function during skin development and inflammation-induced tissue remodelling. Our findings show that podoplanin supports basal layer organisation and the timely acquisition of the epidermal barrier during development, with its loss disrupting cortical organisation and differentiation in keratinocytes. During inflammatory remodelling, conditional deletion alters epidermal differentiation and the response of slow-cycling cells, together with changes in the dermal compartment.

## RESULTS

### Podoplanin expression is tightly regulated in epidermal keratinocytes during morphogenesis in both the interfollicular epidermis (IFE) and hair follicles (HF)

In adult skin, podoplanin is predominantly expressed in lymphatic endothelial cells and is generally absent from epidermal keratinocytes and fibroblasts under physiological conditions, unless upregulated by specific stimuli, most commonly of inflammatory origin (Asai, 2022; Gandarillas et al., 1997). Nevertheless, its spatiotemporal expression during epidermal morphogenesis remains largely uncharacterised. To address this, we investigated the temporal and spatial dynamics of podoplanin expression in the developing epidermis by immunofluorescence and confocal microscopy in mouse samples at distinct embryonic (E12.5, E14.5, E16.5, and E18.5) and postnatal stages (P0, P3, and P5). As shown in Figure 1A, podoplanin was initially expressed in the primitive ectoderm at E12.5-E14.5 and progressively became restricted to the basal layer of the developing epidermis from E16.5 onwards, following a pattern similar to that of keratins 5 and 14 (K5/K14). Notably, regional differences in podoplanin expression varied with developmental stage, with higher levels in the abdominal than in the dorsal epidermis at some stages. This spatiotemporal pattern may reflect regional differences in epidermal maturation, suggesting that podoplanin downregulation occurs later in less mature areas. Furthermore, podoplanin was prominently expressed in other stratified squamous epithelia, including oesophagus and the oral mucosa (Figure S1). In all cases, its expression pattern closely aligned with that of K5/K14, becoming progressively confined to the basal layer of keratinocytes as development proceeded. Importantly, podoplanin expression in epidermal keratinocytes was markedly downregulated during the postnatal period, exhibiting a patchy distribution and becoming detectable only in small clusters of basal epidermal keratinocytes (Figure 1A, arrowheads).

**Figure 1. -.**
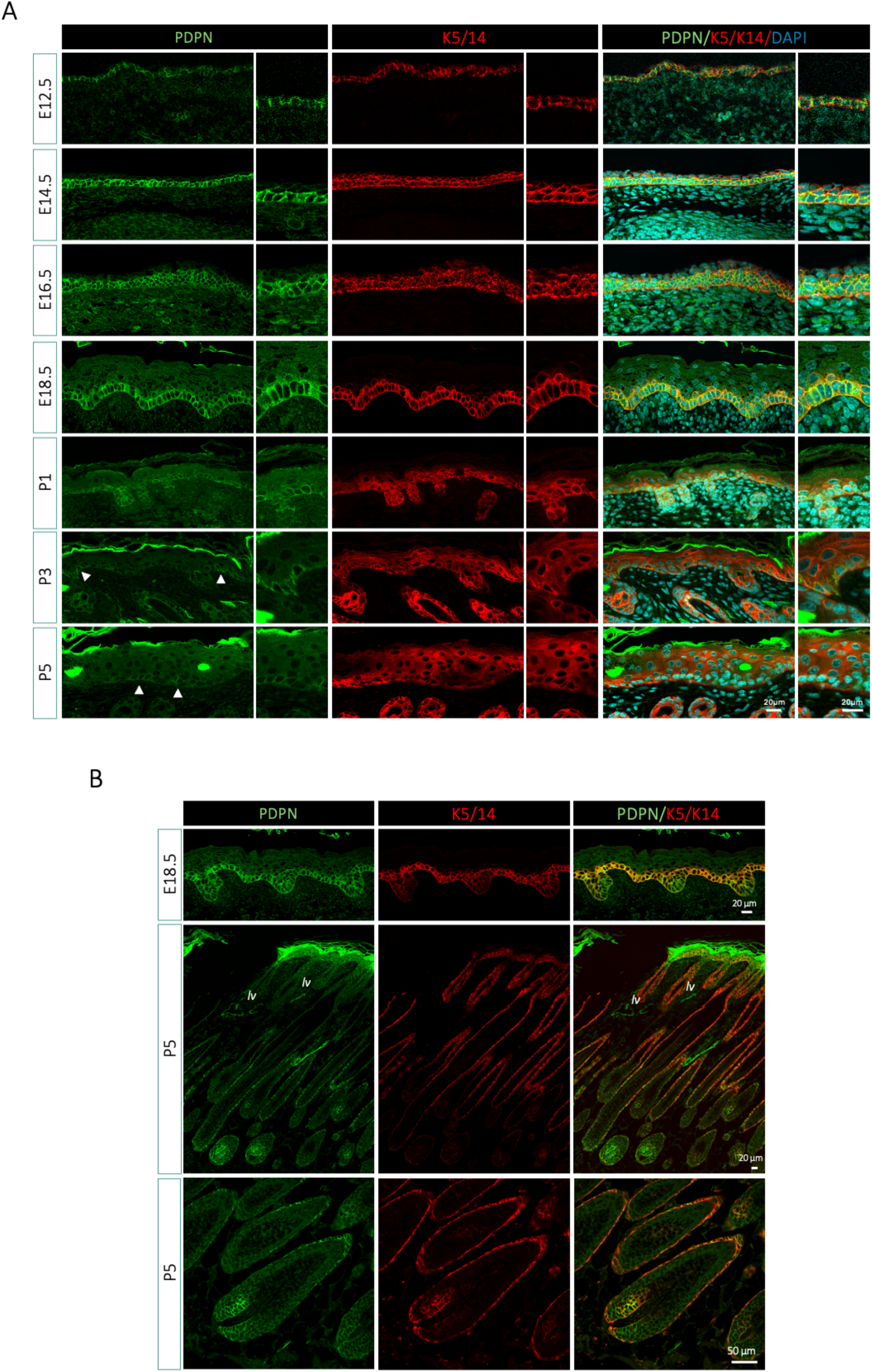
Podoplanin expression during epidermal and hair follicle morphogenesis. *(A)* Representative immunofluorescence images of dorsal skin from wild-type C57BL/6 embryos and neonates at the indicated embryonic (E) and postnatal (P) stages. Sections were stained for podoplanin (PDPN, green) and with a mixture of antibodies against keratins 5 and 14 (K5/K14, red); nuclei were counterstained with DAPI (cyan). Adjacent panels show higher-magnification views. Arrowheads indicate cell clusters retaining podoplanin expression at P3 and P5. *(B)* Representative images highlighting podoplanin expression in developing hair follicles at E18.5 and P5 (morphogenetic anagen). At P5, podoplanin localises to the K5/K14-positive outer root sheath (ORS). Enlarged views of selected areas from the lower-magnification P5 image are shown on the right. Podoplanin-positive lymphatic vessels are indicated (lv). Representative images are shown for each developmental stage (n = 3-4 animals per stage). Scale bars: 20 μm; 50 μm in the enlarged P5 panels.

In addition to its presence in the interfollicular epidermis (IFE), podoplanin was also detected in developing hair follicles (HFs) from E16.5 onwards, eventually becoming restricted to the outer root sheath (ORS) in both primordial and mature HFs (Figure 1B). Notably, in postnatal HFs, podoplanin expression was completely downregulated during the telogen phase and reactivated during the morphogenetic anagen phase (P5), consistent with previous reports (Gandarillas et al., 1997; Yoon et al., 2019). These findings reveal tight spatiotemporal regulation of podoplanin expression during epidermal development and hair follicle cycling. Its enrichment in basal epithelial compartments during tissue growth raises the possibility that podoplanin contributes to the regulation of epidermal stem and progenitor cell behaviour.

### Generation of conditional K5CrePDPN and maternally induced constitutive Pdpn^Δ/Δ^ models

In order to determine the impact of podoplanin expression during epidermal morphogenesis we engineered a mouse model bearing the conditional depletion of podoplanin in the basal layer of the epidermis by crossing *Pdpn*-floxed (Lowe et al., 2015) (*Pdpn*^f/f^) and K5Cre mice (Ramirez et al., 2004) (Figure 2A-D). As shown in Figure 2E, excision of the floxed sequence completely abolished podoplanin expression in primary keratinocytes from newborn mice, as well as in neonatal tissue samples, in a Cre-dependent manner (Figure 2F). As recombination of floxed DNA fragments in K5Cre mice occurs from E15.5 onwards (Ramirez et al., 2004), podoplanin expression was preserved at earlier stages in the basal layer of the epidermis. However, from E16.5 onwards, podoplanin expression started to decline in the epidermis, although residual patchy expression was still observed in occasional basal keratinocytes at E16.5-E18.5. By E18.5, podoplanin was markedly downregulated, albeit not completely abolished, with a more evident persistence in the oral mucosa (Figure S2). This sustained expression suggests both a higher basal expression level of podoplanin in the stratified mucosas compared to the epidermis and an overall high protein stability, indicative of a prolonged half-life in both tissues.

**Figure 2. -.**
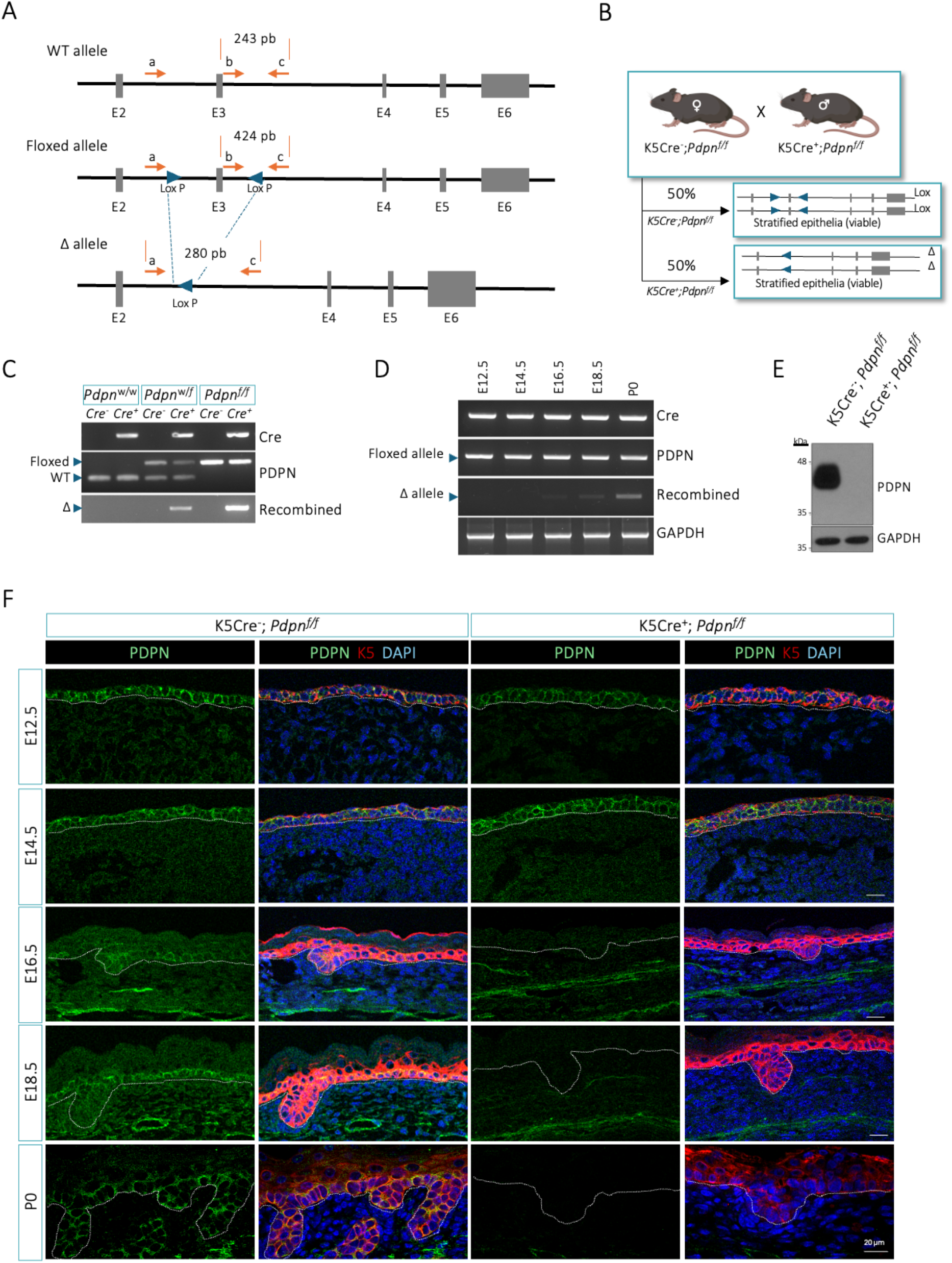
Generation and validation of the epidermal-specific podoplanin knockout model. *(A)* Schematic representation of the wild-type (WT), floxed and recombined Δ *Pdpn* alleles. Exon 3 is flanked by LoxP sites in the floxed allele and excised following Cre-mediated recombination. The positions of primers a, b and c and the expected PCR products for the WT (243 bp), floxed (424 bp) and Δ (280 bp) alleles are indicated. *(B)* Breeding strategy used to generate K5Cre^−^;*Pdpn*^f/f^ control and K5Cre^+^;*Pdpn*^f/f^ conditional KO mice hemizygous for the K5Cre transgene, with an expected frequency of 50% for each genotype. In K5Cre^+^;*Pdpn*^f/f^ mice, Cre-mediated recombination generates the Δ allele in stratified epithelia. *(C)* PCR genotyping of genomic DNA obtained from ear punch biopsies. K5Cre^−^ and K5Cre^+^ samples carrying the indicated *Pdpn*^w/w^, *Pdpn*^w/f^ or *Pdpn*^f/f^ alleles are shown. PCR products corresponding to the Cre transgene and the WT, floxed and Δ *Pdpn* alleles are indicated. *(D)* PCR analysis of genomic DNA obtained from tail tissue at the indicated embryonic and postnatal stages, showing the developmental onset of Cre-mediated recombination in K5Cre^+^;*Pdpn*^f/f^ mice. GAPDH was used as an amplification control. *(E)* Western blot analysis of podoplanin expression in primary keratinocytes isolated from P3-P4 K5Cre^-^;*Pdpn*^f/f^ and K5Cre^+^;*Pdpn*^f/f^ mice. GAPDH was used as a loading control. *(F)* Representative immunofluorescence images of dorsal skin from K5Cre^-^;*Pdpn*^f/f^ and K5Cre^+^;*Pdpn*^f/f^ embryos and neonates at the indicated stages, showing the progressive loss of epidermal podoplanin expression in conditional KO mice. Sections were stained for podoplanin (green) and K5 (red), with DAPI as a nuclear counterstain (blue). White lines delineate the dermo-epidermal junction. Representative images are shown for each developmental stage (n = 3-9 animals per stage). Scale bars, 20 μm

To overcome the delayed loss of podoplanin during development, we took advantage of a distinct feature of the K5Cre model used to generate K5CrePDPN mice. When the K5Cre transgene is maternally inherited, Cre-mediated recombination of floxed alleles can occur in the germline or early embryo and, consequently, throughout all cells and tissues of the progeny, independently of tissue-specific K5 expression and even in offspring that do not inherit the K5Cre transgene (Ramirez et al., 2004). Thus, by modifying our breeding strategy accordingly, we generated mice with constitutive systemic deletion of podoplanin from the onset of embryonic development (Figure 3A-C). Among the resulting progeny, K5Cre^−^;*Pdpn^w/w^* mice were selected as controls and are hereafter referred to as *Pdpn^w/w^*, whereas K5Cre^−^;*Pdpn^Δ/Δ^* mice carrying two recombined *Pdpn* alleles were selected as constitutive KO mice and are hereafter referred to as *Pdpn^Δ/Δ^*. This maternally induced constitutive podoplanin KO model phenocopied the previously described conventional *Pdpn^−/−^* mice, including their perinatal lethality and characteristic lymphatic vascular defects (Cheok et al., 2024; Schacht et al., 2003), enabling us to assess the role of podoplanin in epidermal morphogenesis alongside the K5CrePDPN model. As shown in Figure 3B, recombination of the floxed *Pdpn* alleles occurred even in progeny that did not inherit the K5Cre transgene. Accordingly, podoplanin was absent not only from primary keratinocytes isolated from newborn *Pdpn^Δ/Δ^* mice (Figure 3D), but also throughout the skin and oral mucosa from the onset of embryonic development (Figure 3D and S2C-D).

**Figure 3. -.**
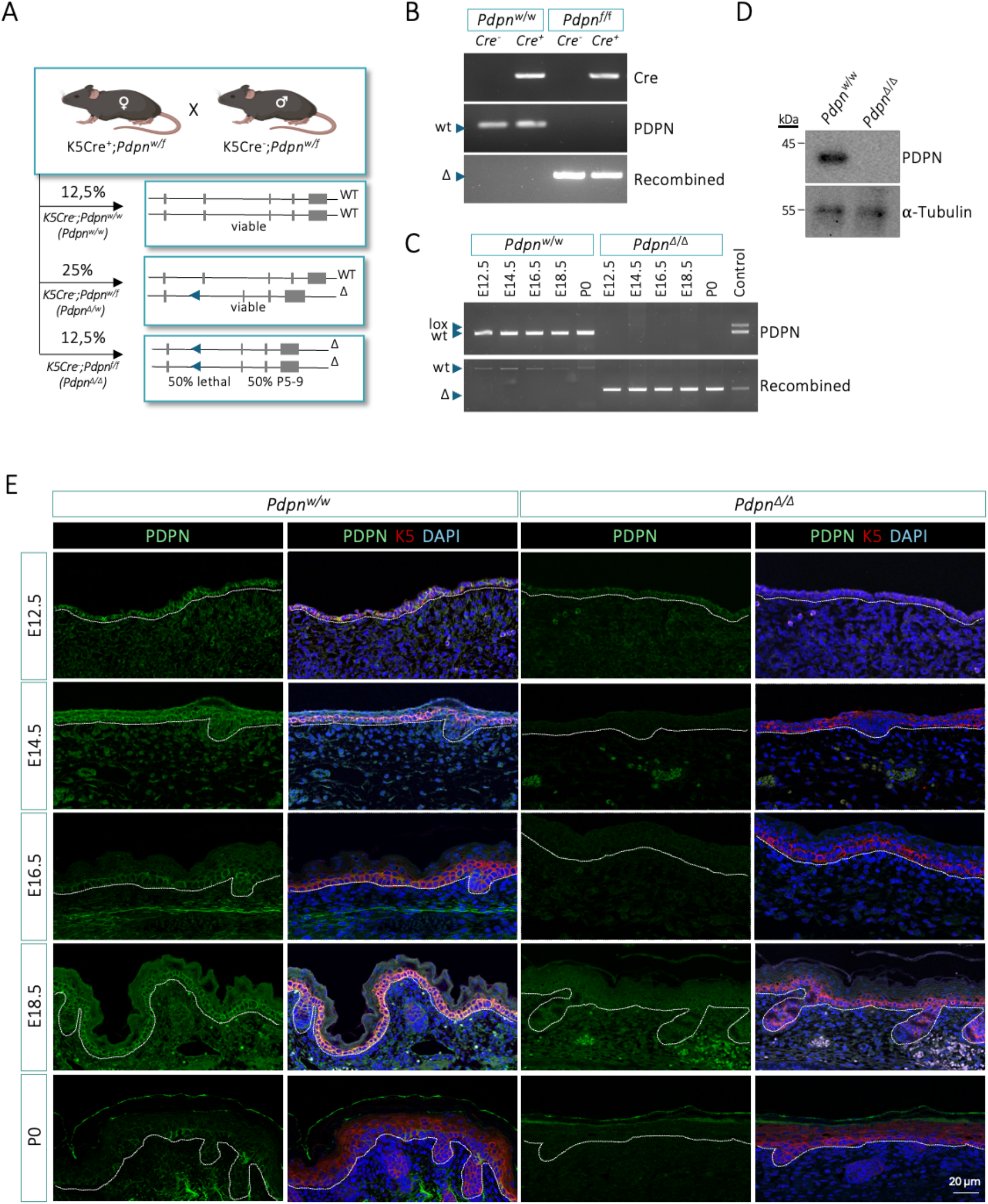
Maternal K5Cre activity results in systemic podoplanin deletion. *(A)* Breeding strategy used to generate constitutive *Pdpn*^Δ/Δ^ mice. K5Cre^+^;*Pdpn*^w/f^ females were crossed with K5Cre^-^;*Pdpn*^w/f^ males. Maternal transmission of K5Cre induced early, widespread recombination of the floxed *Pdpn* alleles, including in offspring that did not inherit the Cre transgene. Only the expected frequencies of the K5Cre^-^ offspring relevant to this study are shown; K5Cre^+^ offspring are omitted. K5Cre^-^;*Pdpn*^w/w^ and K5Cre^-^;*Pdpn*^f/f^ offspring are referred to throughout the manuscript as *Pdpn*^w/w^ and *Pdpn*^Δ/Δ^, respectively. Approximately 50% of *Pdpn*^Δ/Δ^ mice died at birth from respiratory failure, whereas the surviving animals died between P5 and P9. *(B)* PCR analysis of genomic DNA obtained from adult ear punch biopsies. PCR confirmed the presence of the recombined Δ allele in both Cre^−^ and Cre^+^;*Pdpn^f/f^* offspring derived from K5Cre^+^ females. *(C)* PCR analysis of genomic DNA obtained from tail tissue of *Pdpn*^w/w^ and *Pdpn*^Δ/Δ^ embryos and neonates at the indicated stages. Internal positive PCR controls indicate the expected sizes of the WT, floxed and recombined Δ alleles. *(D)* Western blot analysis of podoplanin expression in primary keratinocytes isolated from P3-P4 *Pdpn*^w/w^ and *Pdpn*^Δ/Δ^ mice. α-Tubulin was used as a loading control. *(E)* Representative immunofluorescence images of dorsal skin from *Pdpn*^w/w^ and *Pdpn*^Δ/Δ^ embryos and neonates at the indicated stages, confirming the loss of podoplanin expression throughout development. Sections were stained for podoplanin (green) and K5 (red), with DAPI as a nuclear counterstain (blue). White dotted lines delineate the dermo-epidermal junction. Representative images are shown for each developmental stage (n = 3-9 animals per stage). Scale bar, 20 μm.

### Global and K5Cre-mediated podoplanin deletion reveal a shared epidermal phenotype during skin development

To investigate the consequences of podoplanin loss during skin morphogenesis, we performed a histological time-course analysis of *Pdpn*^Δ/Δ^ and K5CrePDPN mice using haematoxylin and eosin (H&E) staining. Embryonic skin was examined from E12.5 to E18.5 (Figures 4 and 5), followed by dorsal skin at birth (P0) and P5 (Figure 6). The global knockout model allowed us to assess early developmental abnormalities before efficient K5Cre-mediated deletion, while comparison from E16.5 onwards identified epidermal defects shared by both models.

**Figure 4. -.**
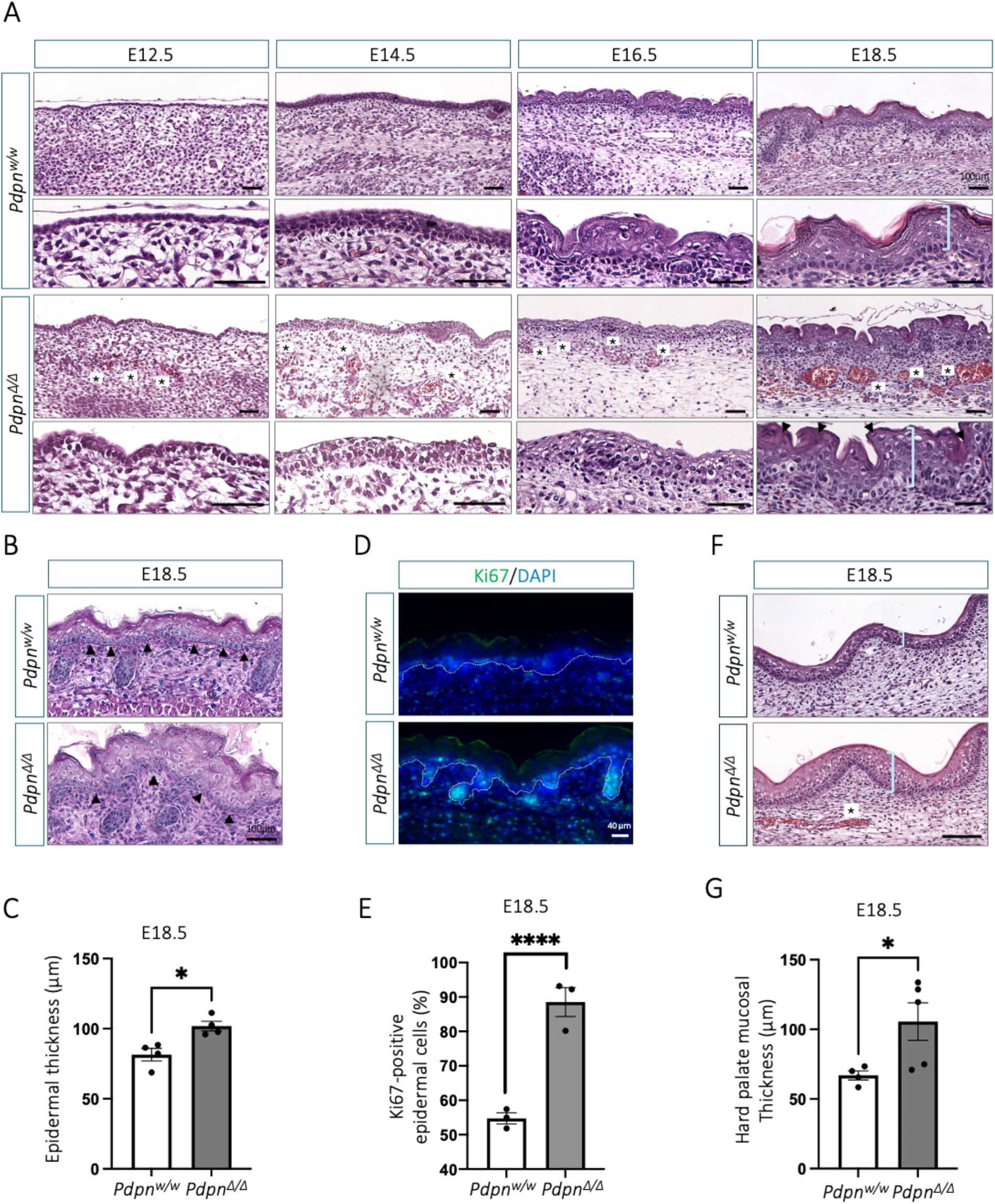
Systemic podoplanin deletion disrupts skin and hard palate morphogenesis. *(A)* Representative haematoxylin and eosin (H&E)-stained sections of dorsal skin from *Pdpn*^w/w^ and *Pdpn*^Δ/Δ^ embryos at the indicated developmental stages (n = 4-15). Lower panels show higher magnification views of the corresponding areas. Asterisks indicate dilated or congested dermal vessels and haemorrhagic foci. Arrowheads at E18.5 indicate retained nuclei in the cornified layer, consistent with parakeratosis. *(B)* Representative PAS-Alcian blue-stained sections of dorsal skin at E18.5, used to visualise the basement membrane and dermo-epidermal junction. Arrowheads highlight the organisation of basal keratinocytes and the basement membrane, which is continuous in *Pdpn^w/w^* skin but shows focal discontinuities in *Pdpn^Δ/Δ^* skin. (n = 4) *(C)* Quantification of epidermal thickness at E18.5 (n = 4 embryos per genotype). *(D)* Representative immunofluorescence images and quantification *(E)* of epidermal proliferation at E18.5. Sections were stained for Ki67 (green), with DAPI as a nuclear counterstain (blue). White dotted lines delineate the dermo-epidermal junction. Ki67-positive cells were quantified as a percentage of the total number of epidermal nuclei (n = 3 embryos per genotype). *(F)* Representative H&E-stained sections of the hard palate from *Pdpn*^w/w^ and *Pdpn*^Δ/Δ^ embryos at E18.5. *(G)* Quantification of total hard palate mucosal thickness at E18.5 (*Pdpn*^w/w^, n = 4; *Pdpn*^Δ/Δ^, n = 5). Brackets indicate epidermal or oral epithelial thickness, as appropriate. Data are presented as mean ± SEM, with individual embryos shown. Statistical significance was determined using two-tailed unpaired Student’s *t*-tests. *p < 0.05; ****p < 0.0001. Scale bars, 100 μm in the H&E and PAS-Alcian blue images and 40 μm in the Ki67 immunofluorescence images.

**Figure 5. -.**
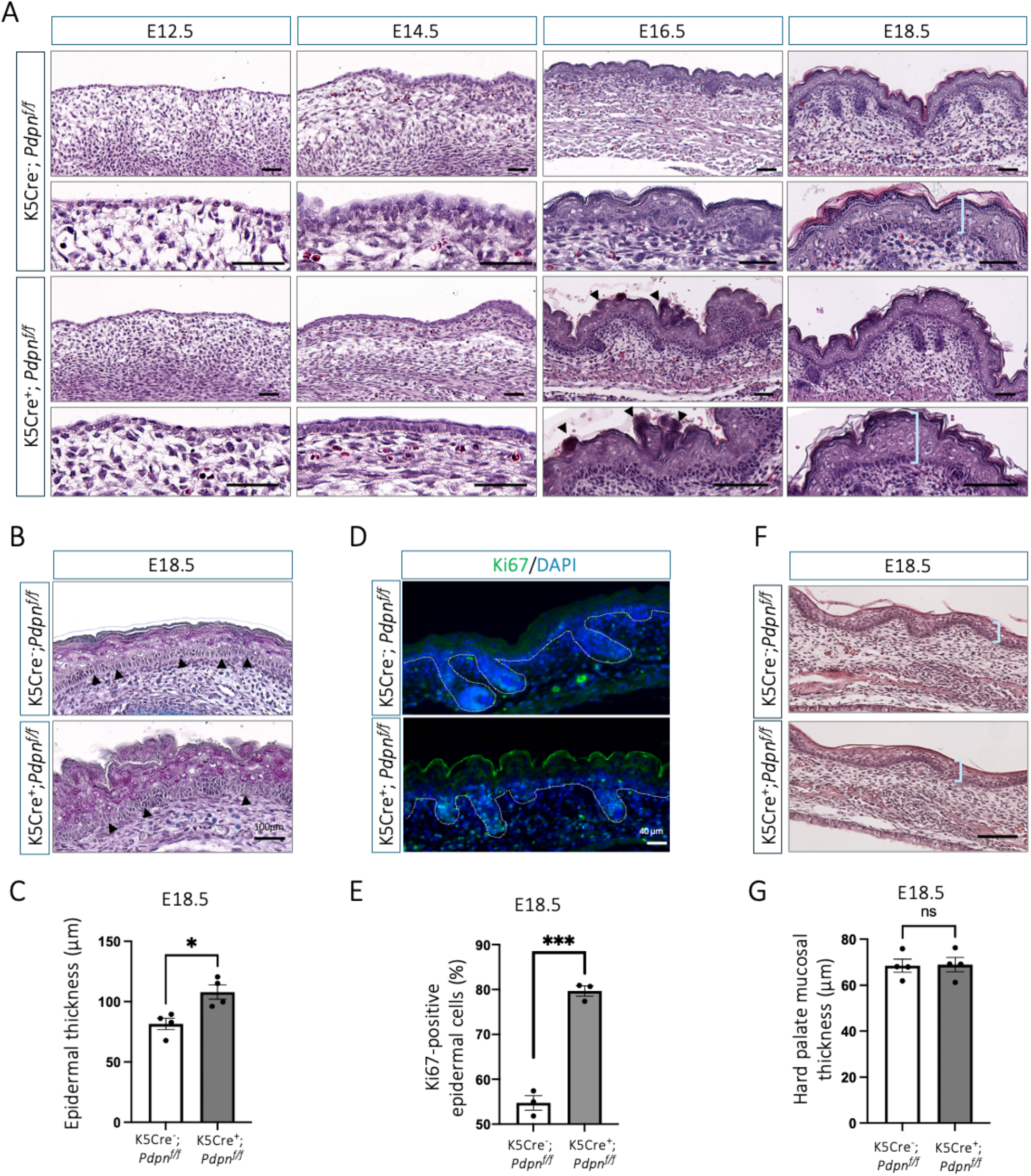
Epidermal-specific podoplanin deletion disrupts skin morphogenesis. *(A)* Representative H&E-stained sections of dorsal skin from K5Cre^-^ ;*Pdpn*^f/f^ and K5Cre^+^;*Pdpn*^f/f^ embryos at the indicated developmental stages. Lower panels show higher-magnification views of the corresponding fields. Arrowheads indicate foci of parakeratosis. (n = 3-9) *(B)* Representative PAS-Alcian blue-stained sections of dorsal skin at E18.5 showing the basement membrane and organisation of basal keratinocytes. Arrowheads highlight the continuous basement membrane in control skin and areas of focal basement membrane disruption in K5Cre^+^;*Pdpn*^f/f^ skin. (n = 4) *(C)* Quantification of epidermal thickness at E18.5 (n = 4 embryos per genotype). *(D)* Representative immunofluorescence images and quantification *(E)* of epidermal proliferation at E18.5. Sections were stained for Ki67 (green) and DAPI as a nuclear counterstain (blue). White dotted lines delineate the dermo-epidermal junction. The percentage of Ki67-positive nuclei was calculated relative to the total number of epidermal nuclei. (n = 3 embryos per genotype). *(F)* Representative H&E-stained sections of the hard palate at E18.5. *(G)* Quantification of total hard palate mucosal thickness at E18.5 (n = 4 embryos per genotype). Brackets indicate epidermal or oral epithelial thickness, as appropriate. Data are presented as mean ± SEM, with individual embryos shown. Statistical analysis was performed using two-tailed unpaired Student’s *t*-tests. *p < 0.05; ***p < 0.001; ns, not significant. Scale bars, 100 μm in H&E and PAS-Alcian blue images and 40 μm in Ki67 immunofluorescence images.

**Figure 6. -.**
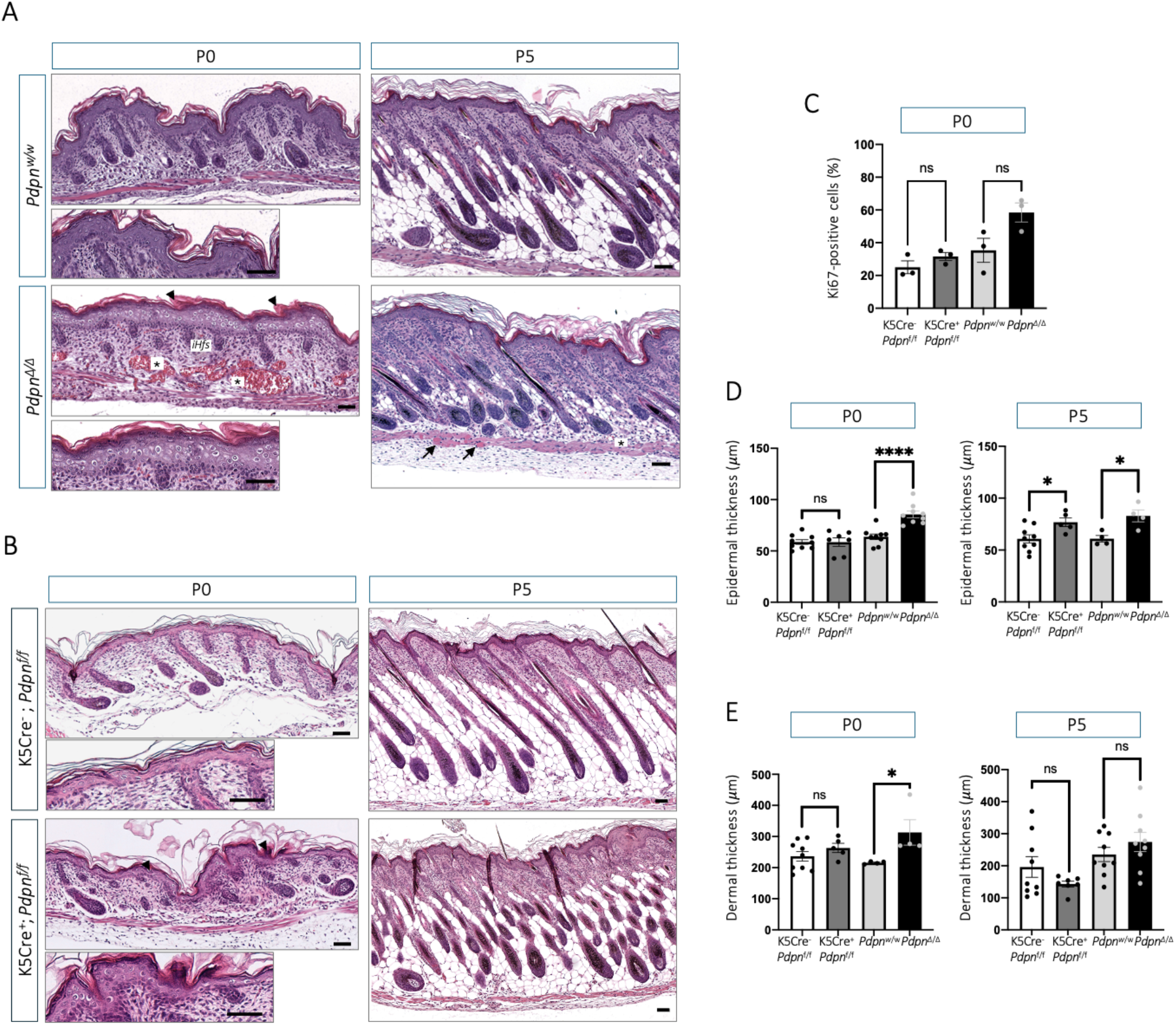
Constitutive and epidermal-specific podoplanin KO models exhibit overlapping neonatal epidermal phenotypes. *(A)* Representative H&E-stained sections of dorsal skin from *Pdpn*^w/w^ and *Pdpn*^Δ/Δ^ neonates at P0 (n = 9 animals per genotype) and P5 (n = 4-15 animals per genotype). Lower P0 panels show higher-magnification views of the corresponding fields. Arrowheads indicate parakeratotic foci, asterisks indicate vascular congestion and arrows at P5 indicate focal hyaline degeneration in the underlying muscle. iHfs; immature hair follicles. *(B)* Representative H&E-stained sections of dorsal skin from K5Cre^-^;*Pdpn^f/f^* and K5Cre^+^;*Pdpn^f/f^* neonates at P0 (n = 7-9 animals per genotype) and P5 (n = 5-10 animals per genotype). Lower P0 panels show higher-magnification views. Arrowheads indicate parakeratotic foci. *(C)* Quantification of Ki67-positive nuclei relative to the total number of epidermal nuclei at P0, (n = 3 animals per group). *(D)* Quantification of epidermal (D) and dermal (E) thickness at P0 and P5. Data are presented as mean ± SEM, with individual animals shown (n = 3-9 animals per group; exact n values correspond to the number of plotted points). Comparisons were performed separately between control and knockout animals within each model and age using two-tailed unpaired Student’s t-tests. *p < 0.05; ****p < 0.0001; ns, not significant. Scale bars, 100 μm.

In *Pdpn*^Δ/Δ^ embryos, early epidermal abnormalities included focal disruption of basal keratinocyte nuclear polarity and cytoplasmic vacuolisation, consistent with intracellular oedema. These changes became more pronounced between E14.5 and E16.5, a period of active epidermal stratification, and were accompanied by focal epidermal thickening (Figure 4A). K5CrePDPN embryos showed no detectable histological abnormalities at E12.5 or E14.5, consistent with the later onset of efficient podoplanin deletion (Ramirez et al., 2004) (Figure 5A). By E16.5, however, both models displayed focal basal layer disorganisation and keratinocyte vacuolisation, with greater severity in *Pdpn*^Δ/Δ^ skin (Figures 4A and 5A). At E18.5, basal and suprabasal vacuolisation persisted alongside basal disorganisation, and abnormalities in epidermal keratinisation, including focal parakeratosis (Figures 4A and 5A, arrowheads). Focal basement membrane disruption was also detected in both models, particularly in embryos with more severe epidermal abnormalities (Figures 4B and 5B, arrowheads). These alterations in epidermal architecture and differentiation were accompanied by increased keratinocyte proliferation and epidermal thickness relative to the respective controls (Figures 4C-D and 5C-D). In addition, histological alterations also affected the oral mucosa in *Pdpn*^Δ/Δ^ mice, including epithelial thickening, intracellular oedema of keratinocytes and vascular congestion in the underlying submucosa, whereas no significant increase in oral mucosal thickness was detected in K5CrePDPN mice (Figures 4E-F and 5E-F).

Several epidermal abnormalities persisted after birth. At P0, *Pdpn*^Δ/Δ^ skin retained suprabasal vacuolisation, focal basal nuclear disorganisation and a thickened cornified layer with retained nuclei, consistent with parakeratosis (Figure 6A, arrowheads). The epidermal surface was generally smooth, with less prominent dermo-epidermal ridges, although ridges remained evident in some animals. K5CrePDPN skin showed more focal abnormalities, including epidermal hyperplasia and parakeratosis (Figure 6B). At this stage, the increase in keratinocyte proliferation was no longer significant in either model (Figure 6C). Epidermal thickness nevertheless remained significantly increased in *Pdpn*^Δ/Δ^ mice, while focal hyperplasia in K5CrePDPN skin did not translate into a significant overall increase. Hair follicle differentiation was also delayed in *Pdpn*^Δ/Δ^ skin at P0, with follicles remaining at earlier developmental stages than those in *Pdpn*^w/w^ controls (Figure 6A). These follicles were shorter, located in the superficial dermis and lacked a clearly differentiated dermal papilla within the hair bulb, whereas control follicles had extended into the deep dermis, close to the subcutaneous muscle, and displayed more advanced hair bulb differentiation. Alterations in follicular development were also observed in K5CrePDPN skin. By P5, follicles in both mouse models had generally progressed into anagen and epidermal thickness was significantly increased in both models (Figures 6A-B).

The global KO additionally displayed marked vascular and interstitial abnormalities. Mild interstitial oedema and dilation of dermal vessels were detectable at E12.5, increased from E14.5 and were most pronounced at E16.5 and E18.5, when vascular congestion, microhaemorrhages and marked lymphatic dilation were evident (Figure 4A, asterisks). These findings recapitulated the phenotype reported in conventional *Pdpn*^-/-^ mice on a 129/Sv background (Schacht et al., 2003). Oedema varied in extent and distribution among embryos: diffuse oedema was generally associated with a thin, flattened epidermis, whereas focal oedema was associated with prominent dermo-epidermal ridges and severe vessel congestion. Dermal fibroblasts also displayed cytoplasmic vacuolisation consistent with intracellular oedema. Postnatally, abnormalities in deeper skin compartments were mainly observed in *Pdpn*^Δ/Δ^ mice, including small, poorly developed hypodermal adipocytes and focal hyaline degeneration of muscle fibres at P5 (Figure 6A, arrows). Dermal thickness was significantly increased only in *Pdpn*^Δ/Δ^ mice at P0, with no significant difference at P5 (Figure 6E). Increased dermal cellularity, however, was observed in both models, showing that changes accompanying podoplanin deletion in basal keratinocytes extended beyond the epidermal compartment.

Together, these findings demonstrate that podoplanin deletion in basal keratinocytes disrupts epidermal organisation and follicular differentiation during skin development and is accompanied by increased dermal cellularity. Global deletion produces a more severe cutaneous phenotype, with additional vascular and interstitial abnormalities. The shared epidermal defects establish a contribution of podoplanin in keratinocytes to skin morphogenesis, while the dermal changes in the conditional model indicate that its effects extend beyond the epithelial compartment.

### Podoplanin loss delays skin barrier development and drives imbalanced epidermal differentiation in vivo

To investigate whether the observed phenotypic changes were linked to functional skin defects, we evaluated epidermal barrier formation in both mouse models using a toluidine blue-based dye permeability assay. Barrier formation typically initiates dorsally and progressively extends to the ventral surface, starting at E16.5 and becoming fully established by E18.5 (Hardman et al., 1998). In *Pdpn^Δ/Δ^* mice, delayed barrier formation was evident at E16.5 compared to *Pdpn^w/w^*controls, with the main differences persisting at E17.5 in ventral and lateral skin regions (Figure 7A-B). By E18.5, the barrier was fully restored, suggesting the activation of compensatory mechanisms. Similarly, conditional podoplanin KO mice exhibited a comparable delay. At E16.5, however, barrier formation was not yet evident in either conditional KO embryos or their corresponding controls. Accordingly, significant defects were only detected at E17.5, mainly affecting the lateral and ventral skin (Figure 7C and D).

**Figure 7. -.**
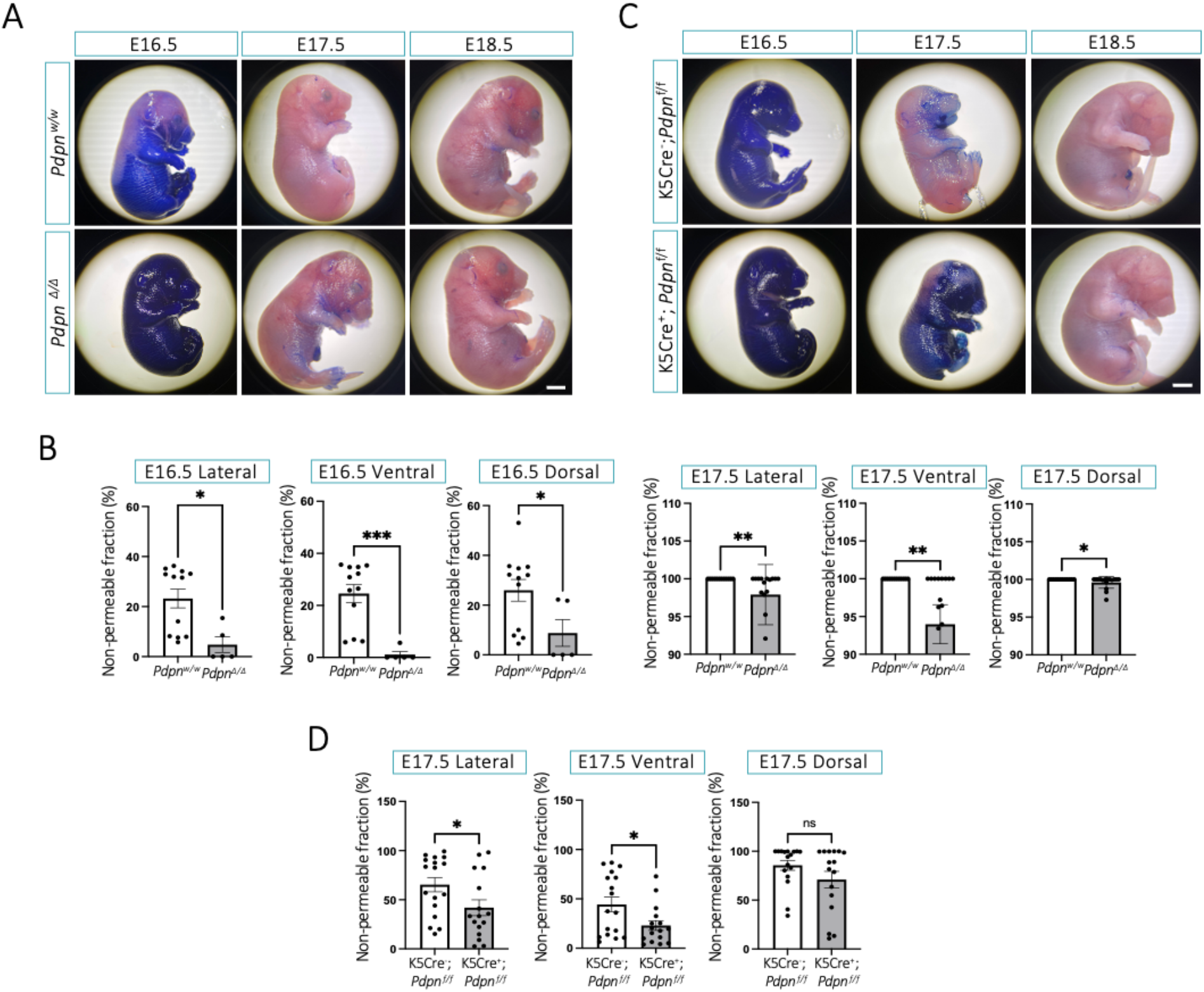
Podoplanin deficiency delays epidermal barrier acquisition during embryonic development. *(A)* Representative lateral views of *Pdpn*^w/w^ and *Pdpn*^Δ/Δ^ embryos at E16.5, E17.5 and E18.5 following the toluidine blue permeability assay. Toluidine blue penetration indicates incomplete epidermal barrier formation, whereas dye exclusion indicates the acquisition of a functional barrier. *(B)* Quantification of the non-permeable surface fraction in lateral, ventral and dorsal views of *Pdpn*^w/w^ and *Pdpn*^Δ/Δ^ embryos at E16.5 and E17.5. *(C)* Representative lateral views of K5Cre^-^;*Pdpn^f/f^* and K5Cre^+^;*Pdpn^f/f^* embryos at the indicated developmental stages. *(D)* Quantification of the non-permeable surface fraction in the three orientations of K5Cre^-^;*Pdpn^f/f^* and K5Cre^+^;*Pdpn^f/f^* embryos at E17.5. Data are presented as mean ± SEM, with each point representing an individual embryo (*n* = 5-17 embryos per group; exact sample sizes are indicated by the individual data points). Genotypes were compared separately within each developmental stage and orientation using two-tailed unpaired Student’s *t*-tests. ns, not significant;\**P* < 0.05, **P < 0.01 and \*\*\**P* < 0.001. Scale bars, 1mm.

To further characterise epidermal differentiation during this developmental window, we analysed the expression of basal and differentiation markers by immunofluorescence at late embryonic and early postnatal stages. In the global KO model, *Pdpn^Δ/Δ^* embryos were analysed at E16.5, E18.5 and P0 (Figure 8A), whereas conditional KO embryos were examined at E17.5, E18.5 and P0 (Figure 8B). In *Pdpn^Δ/Δ^* embryos, K5 staining displayed a patchy and discontinuous pattern compared to *Pdpn^w/w^* controls, consistent with altered basal layer organisation and irregular keratinocyte morphology. A similar, disruption in K5 distribution was also observed in conditional KO embryos. Both KO models displayed an increased number of K10-positive suprabasal keratinocyte layers, consistent with the epidermal hyperplasia observed histologically. In addition, aberrant K10 expression in superficial parakeratotic layers, was also focally observed (Figure 8A, E18.5), indicating abnormal epidermal differentiation and keratinisation. Despite the delay in barrier acquisition described above, both mouse models also displayed earlier, more prominent loricrin staining, indicating premature expression of a terminal differentiation marker.

**Figure 8. -.**
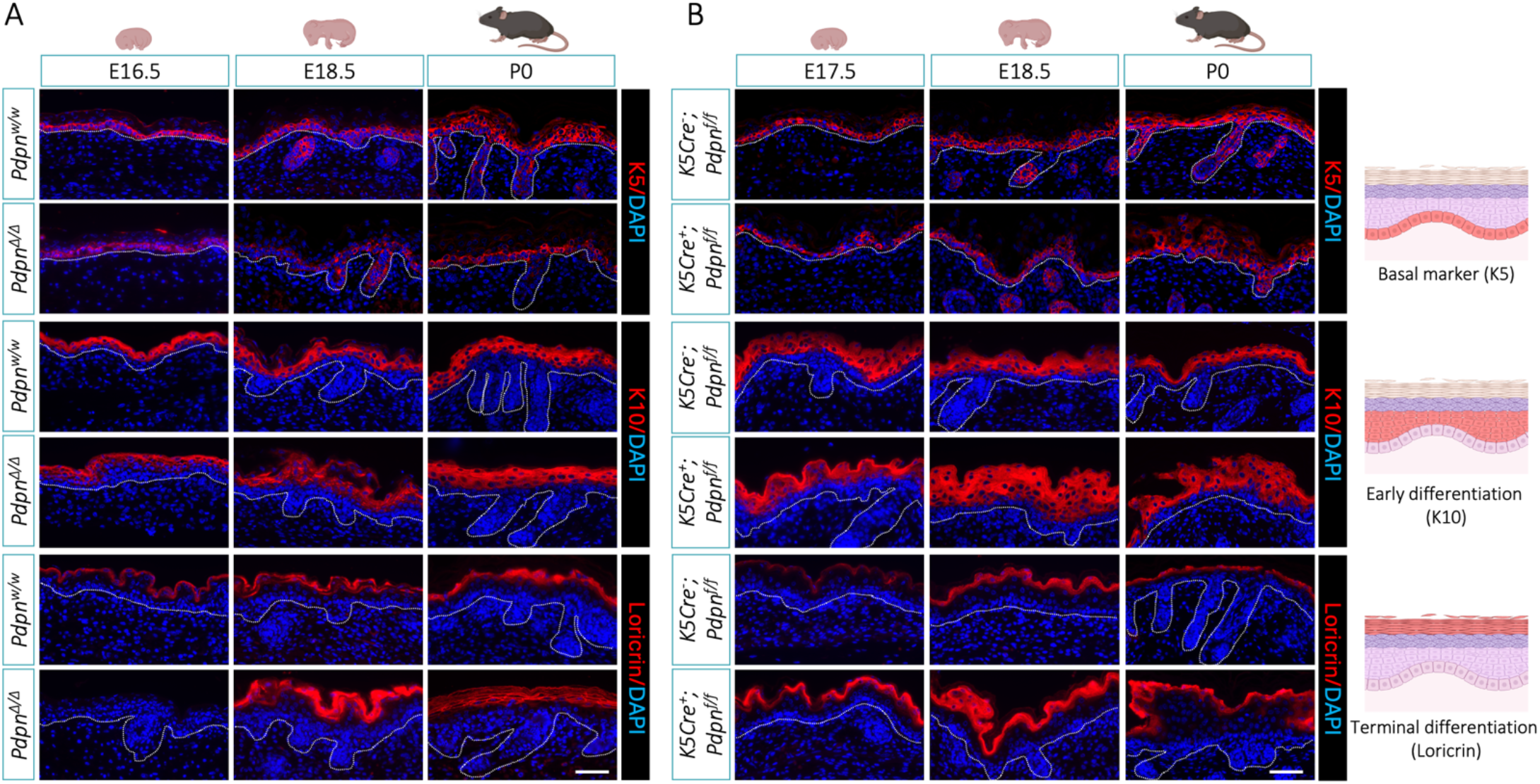
Podoplanin loss disrupts epidermal differentiation during late embryonic and early postnatal development. *(A)* Representative immunofluorescence images of dorsal skin from *Pdpn*^w/w^ and *Pdpn*^Δ/Δ^ mice at E16.5, E18.5 and P0. *(B)* Representative immunofluorescence images of dorsal skin from K5Cre^-^;*Pdpn^f/f^* and K5Cre^+^;*Pdpn^f/f^* mice at E17.5, E18.5 and P0. Sections were stained for K5, K10 or loricrin (red), with DAPI nuclear counterstaining (blue). White dotted lines indicate the epidermal–dermal boundary. The images illustrate alterations in the basal restriction of K5, expansion of suprabasal K10 expression and changes in the spatiotemporal expression pattern of loricrin following podoplanin loss. Scale bars, 50 µm.

Together, these results show that podoplanin deficiency uncouples the expression of differentiation markers from timely acquisition of epidermal barrier function. The expanded and aberrant distribution of K10, together with premature loricrin expression, was accompanied by basal layer disorganisation, cornified layer thickening and focal parakeratosis, indicating that altered differentiation marker expression did not translate into faster functional maturation. Thus, podoplanin contributes to the coordination of epidermal differentiation and barrier acquisition during skin morphogenesis.

### Podoplanin deficiency disrupts basal keratinocyte division orientation during epidermal morphogenesis

Epidermal morphogenesis requires coordination between expansion of the basal progenitor compartment and formation of the suprabasal layers. Basal keratinocyte division orientation is shaped by the interplay between proliferation-induced crowding, cell geometry and adhesion, together with apico-basal polarity and cortical cues that position the mitotic spindle (Jiang et al., 2021), (Box et al., 2019; Luxenburg et al., 2015). Given our findings of basal layer disorganisation, altered stratification and an imbalance between proliferation and differentiation following podoplanin loss, we next examined whether basal keratinocyte division orientation was also affected in *Pdpn*Δ/Δ and K5CrePDPN mice. During epidermal development, planar (symmetric) divisions generally retain both daughter cells within the basal layer, thereby contributing to expansion of the basal progenitor compartment, whereas perpendicular (asymmetric) divisions can place one daughter in a suprabasal position and have classically been linked to epidermal stratification and differentiation (Gadre et al., 2024; Williams et al., 2011). We therefore quantified basal keratinocyte division orientation using phospho-histone H3 (pHH3) immunostaining together with the morphology of DAPI-labelled chromatin to identify mitotic figures. Only pHH3-positive basal cells displaying two clearly separated chromatin masses were included, and division orientation was determined by measuring the angle between the axis connecting the centres of these masses and the local epidermal-dermal interface. Divisions were classified as planar when occurring at angles below 30°, and as perpendicular when exceeding 60°, while intermediate angles (30-60°) were considered oblique (Figure 9A). To examine the developmental redistribution of basal keratinocyte division orientations, analyses were performed at E12.5 and during late embryonic development (E16.5-E18.5), when the frequency of perpendicular divisions normally increases.

**Figure 9. -.**
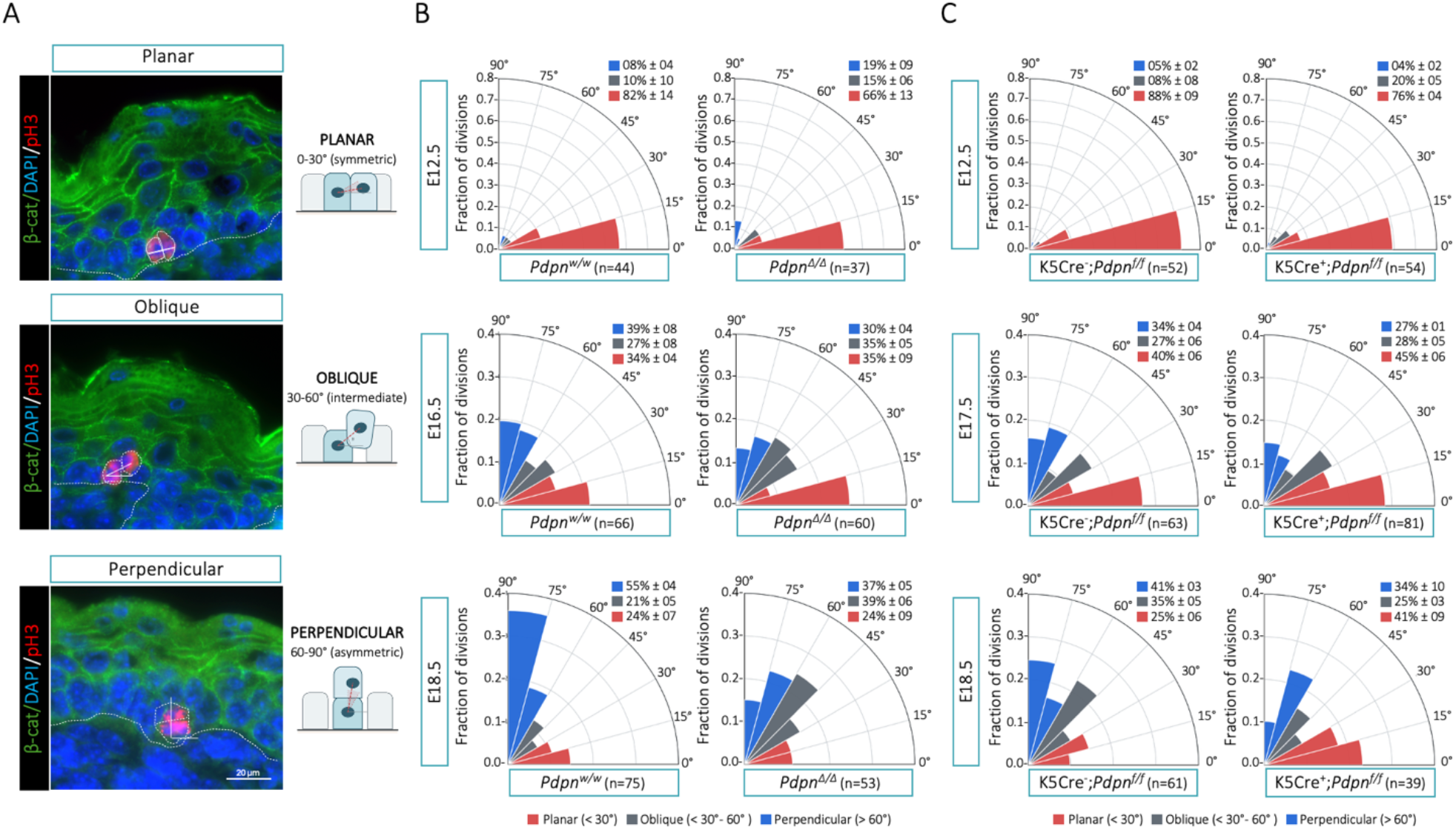
Podoplanin loss alters the orientation of basal epidermal cell divisions during skin morphogenesis. *(A)* Representative immunofluorescence images of planar, oblique and perpendicular mitotic divisions in basal epidermal cells. Sections were stained for β-catenin (green) to visualise cell boundaries, phospho-histone H3 (pHH3; red) to identify mitotic cells and DAPI (blue) to label nuclei. The dashed white line indicates the basement membrane, and the solid lines illustrate the mitotic axis and its orientation relative to the basement membrane. Mitotic divisions were classified as planar (0-30°), oblique (30-60°) or perpendicular (60-90°). *(B)* Polar plots showing the distribution of mitotic division orientations in *Pdpn*^w/w^ and *Pdpn*^Δ/Δ^ epidermis at E12.5, E16.5 and E18.5. *(C)* Distribution of mitotic division orientations in K5Cre^-^;*Pdpn^f/f^* and K5Cre^+^;*Pdpn^f/f^* epidermis at E12.5, E17 and E18.5. Percentages are presented as mean ± SEM from three independent embryos per condition. The values of *n* shown beneath each plot indicate the total number of mitoses analysed. (n = 3 embryos per genotype). Scale bar, 20 µm.

As shown in Figure 9B, both control groups (*Pdpn*^w/w^ and K5Cre^-^;*Pdpn^f/f^*) displayed the expected developmental reorientation of basal keratinocyte divisions, characterised by a sharp increase in perpendicular divisions during late gestation. In *Pdpn^w/w^* embryos, perpendicular divisions increased abruptly at E16.5, rising approximately five-fold compared to E12.5 (8-39%) and reached a ∼7-fold increase by E18.5 (8-55%). A comparable dynamic was observed in Cre^-^;*Pdpn^f^*^/f^ controls, where perpendicular divisions progressively increased at E17.5 and reached an ∼8-fold enrichment by E18.5 (5-34-41%). This transition was accompanied by a marked decline in planar divisions and a transient increase in oblique orientations, consistent with progressive epidermal stratification. In both podoplanin-deficient models, however, the developmental redistribution from planar towards perpendicular division orientations was attenuated, although this alteration manifested differently in each model. In *Pdpn^Δ/Δ^*epidermis, the proportion of perpendicular divisions increased only ∼2-fold between E12.5 and E18.5 (from 19% to 37%) and remained lower than in controls at E18.5 (37% versus 55%). At this stage, the lower proportion of perpendicular divisions was accompanied by an increased proportion of oblique orientations. In K5CrePDPN embryos, oblique divisions were already more frequent at E12.5 than in controls (20% versus 8%), despite incomplete podoplanin depletion, accompanied by a reduction in planar divisions (76% versus 88%). The main difference emerged at later stages, when mutant epidermis failed to show the continued shift towards perpendicular divisions observed in controls. Thus, planar divisions remained frequent in K5Cre^+^;*Pdpn^ᶠ/ᶠ^*embryos at both E17.5 and E18.5 (45% and 41%, respectively), whereas their proportion in K5Cre^-^;*Pdpn^ᶠ/ᶠ^* controls declined from 40% at E17.5 to 25% at E18.5 (Figure 9C).

Overall, these findings show that podoplanin facilitates the timely developmental transition from planar to perpendicular divisions in basal keratinocytes during epidermal morphogenesis. In *PdpnΔ*/*Δ* embryos, this transition was altered by an increased proportion of oblique divisions, whereas in conditional K5Cre+;Pdpnf/f embryos it is characterised by a sustained persistence of planar divisions at late stages. Together, these findings show that podoplanin loss disrupts the developmental control of basal keratinocyte division orientation during epidermal morphogenesis.

### Podoplanin-deficient keratinocytes exhibit an altered temporal response to Ca^2+^-induced differentiation

To determine whether the altered differentiation programme observed *in vivo* reflected a keratinocyte-intrinsic defect, we examined Ca^2+^-induced differentiation in primary keratinocytes isolated from control and mutant K5CrePDPN mice. Differentiation was induced with 1.2 mM CaCl₂, and cultures were analysed at 0, 48 and 120h after stimulation. As shown in Figure 10A, phase-contrast microscopy revealed the expected Ca^2+^-induced morphological response in both genotypes, including increased cell size, flattening, acquisition of a polygonal morphology and enhanced cell-cell cohesion. By 48h, however, podoplanin-deficient keratinocytes displayed more pronounced flattening and multilayering than control cells, consistent with an earlier and more pronounced morphological response to Ca^2+^-induced differentiation. We next examined the expression of epidermal differentiation markers by immunoblotting before 1.2 mM CaCl₂ addition and after 48 and 120 h of stimulation (Figure 10B). In control keratinocytes, Ca^2+^ stimulation induced involucrin and loricrin expression and a progressive reduction in K5, particularly at 120 h, whereas K10 remained weakly expressed. Podoplanin-deficient keratinocytes displayed higher basal levels of K10, involucrin and loricrin than controls, but these markers followed distinct trajectories after Ca^2+^addition. K10 declined by 120 h, while loricrin failed to show the late increase observed in controls and was lower in mutant cells at this time point. Involucrin remained relatively stable between 0 and 48 h before increasing markedly at 120 h. K5 levels were also modestly lower in podoplanin-deficient cells. Thus, the higher basal expression of differentiation markers in podoplanin-deficient cells was not accompanied by a uniformly accelerated response to Ca^2+^, indicating altered temporal coordination of differentiation. Interestingly, in agreement with previous studies in human keratinocytes (Honma et al., 2019), podoplanin itself was consistently downregulated during differentiation in control cells, further supporting its association with an undifferentiated basal state. Together, these findings support a keratinocyte-intrinsic role for podoplanin in coordinating the onset and progression of epidermal differentiation and are consistent with the altered differentiation dynamics observed *in vivo*.

**Figure 10. -.**
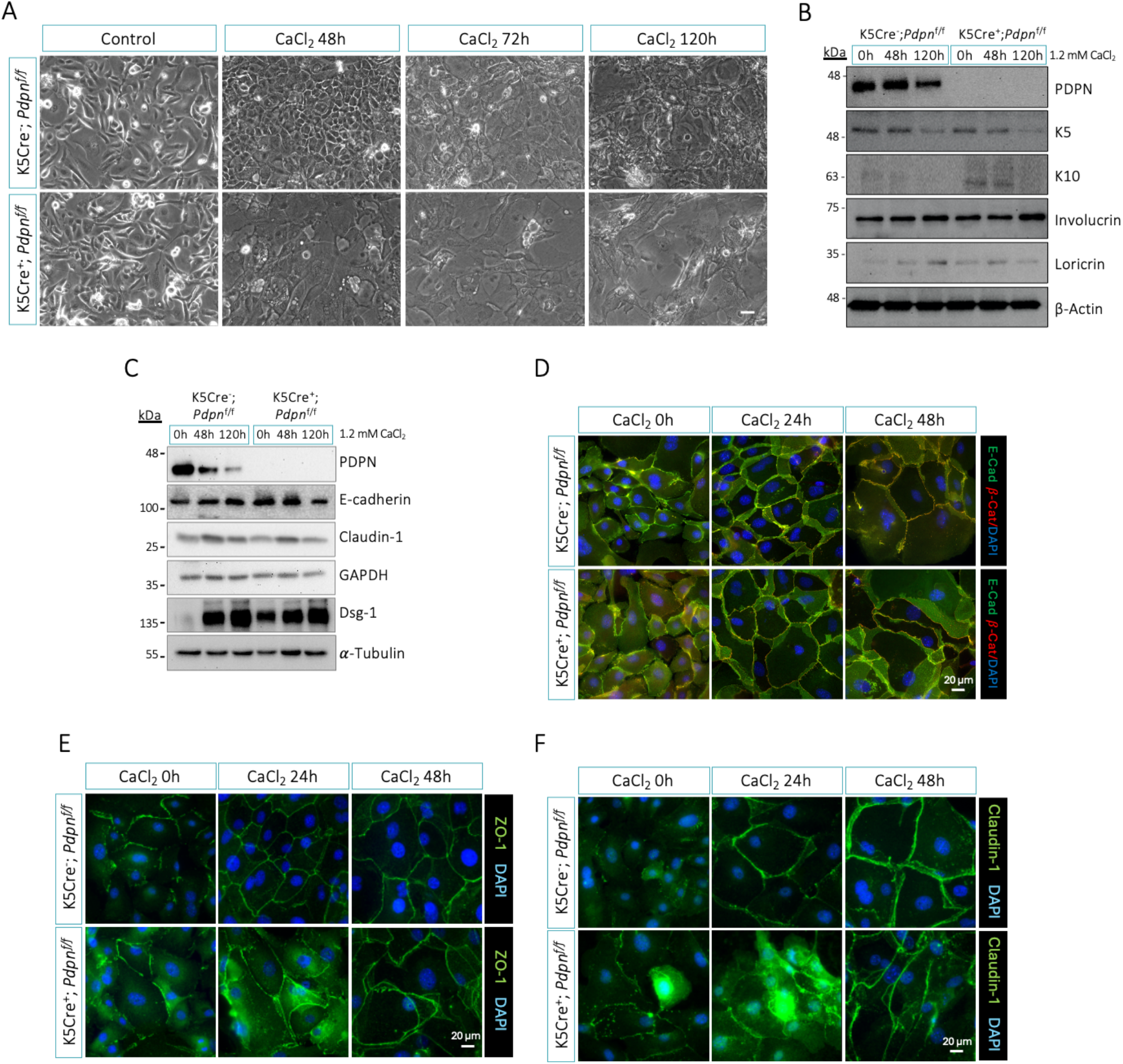
Podoplanin loss alters Ca^2+^-induced differentiation and junctional organisation in primary keratinocytes. Primary keratinocytes isolated from K5Cre^-^;*Pdpn^f/f^* and K5Cre^+^;*Pdpn^f/f^* mice were induced to differentiate by supplementation with 1.2 mM CaCl₂. *(A)* Representative phase-contrast images of keratinocyte cultures before Ca^2+^ addition and after 48, 72 and 120 h of differentiation. *(B)* Western blot analysis of podoplanin and the differentiation markers K5, K10, involucrin and loricrin at 0, 48 and 120 h. β-Actin was used as a loading control. *(C)* Western blot analysis of podoplanin and the junction-associated proteins E-cadherin, claudin-1 and desmoglein-1 (Dsg-1). GAPDH and α-tubulin were used as a loading control. (D) Representative immunofluorescence images showing the distribution of E-cadherin (green) and β-catenin (red) at 0, 24 and 48 h after Ca^2+^ addition. (E– F) Representative immunofluorescence images showing the localisation of ZO-1 (E) and claudin-1 (F) at the indicated time points. Nuclei were counterstained with DAPI (blue). Images and immunoblots are representative of more than three independent keratinocyte isolations. Representative images from at least 4 independent experiments are shown. Scale bars, 50 µm in (A) and 20 µm in (D–F).

### Ca^2+^-induced differentiation of podoplanin-deficient keratinocytes reveals defective cortical actin and cell-cell junction organisation

Given the basal layer disorganisation and altered division orientation observed *in* vivo, together with the previously reported inverse association between podoplanin and E-cadherin expression in mouse skin SCCs (Scholl et al., 1999) and human oral SCCs (Martín-Villar et al., 2005), we next investigated whether podoplanin loss altered the expression and localisation of junctional proteins during Ca^2+^-induced differentiation in primary keratinocytes. As shown in Figure 10C, control keratinocytes elicited the expected modulation of adhesion molecules upon Ca^2+^stimulation over time, consistent with the progressive establishment of epithelial junctions. In contrast, K5Cre^+^;*Pdpn^f/f^*cells exhibited an altered regulatory profile. E-cadherin levels were upregulated under basal conditions in K5Cre^+^;*Pdpn^f/f^* cells but declined at later stages of differentiation, rather than maintaining stable expression or being upregulated. Claudin-1 showed a transient increase at 48 hours following Ca^2+^addition in mutant keratinocytes, which was not sustained at 120 hours. Moreover, desmoglein-1 was also upregulated in podoplanin-deficient keratinocytes under basal conditions although showed the differentiation-associated modulation observed in control cells. Immunofluorescence analysis of the adherens junction proteins E-cadherin and β-catenin and the tight junction proteins ZO-1 and claudin-1 at 24 and 48 h after Ca^2+^addition further revealed altered epithelial organisation in podoplanin-deficient keratinocytes. (Figure 10D-F). ZO-1 and claudin-1 showed greater cytoplasmic accumulation in mutant cells at both 0 and 24 h compared to control cells. Although junctional proteins were still largely recruited to cell-cell contacts following Ca^2+^ stimulation, mutant cells exhibited markedly irregular boundaries and lacked the regular polygonal organisation observed in control monolayers. Together with the altered expression dynamics detected by immunoblotting, these findings indicate that podoplanin loss affects both the temporal regulation of junctional protein abundance and the subcellular distribution of selected components during keratinocyte differentiation.

The alterations in cell geometry prompted us to examine the cortical actomyosin network that mechanically supports epithelial junctions. The rationale for this analysis was further supported by the established interaction of podoplanin with ERM (ezrin, radixin and moesin) proteins and its ability to regulate RhoA/C-dependent actin remodelling in epithelial and SCC cells (Martín-Villar et al., 2015, 2006), as well as actomyosin contractility in fibroblastic reticular cells (Acton et al., 2014; Astarita et al., 2015). Accordingly, we examined ERM and MLC2 phosphorylation and subcellular localisation, together with actin cytoskeleton organisation, during Ca^2+^-induced differentiation. ERM activation was assessed by phosphorylation of ezrin at Thr567, radixin at Thr564 and moesin at Thr558 (collectively referred to as pERM), whereas MLC2 activation was assessed by dual phosphorylation at Thr18 and Ser19 (ppMLC2). As shown in Figure 11A, immunoblot analysis of unstimulated cultures revealed no evident difference in total ERM levels between genotypes, whereas pERM levels were reduced in podoplanin-deficient keratinocytes, indicating decreased basal ERM phosphorylation. By contrast, total MLC2 and ppMLC2 levels were increased to a similar extent in mutant cells, resulting in no evident change in the ppMLC2/MLC2 ratio. Thus, podoplanin loss reduced basal pERM activation while increasing the overall abundance of MLC2 without altering its relative phosphorylation. We next examined the subcellular localisation of pERM and ppMLC2 during early Ca^2+^-induced differentiation (24-48h; Figure11B-C). In unstimulated control keratinocytes, pERM was predominantly detected in cell protrusions and peripheral membrane structures. Following Ca^2+^ addition, pERM progressively accumulated at cell-cell contacts, with junctional enrichment evident at 24 and 48 h. Podoplanin-deficient keratinocytes displayed weaker pERM staining before stimulation and did not show comparable junctional enrichment following Ca^2+^ treatment (Figure11B). Similarly, ppMLC2 was associated with stress fibres and discrete cortical regions in unstimulated control cells. Ca^2+^ stimulation induced its progressive redistribution towards the cortical cytoskeleton at cell-cell contacts, which was more pronounced at 48 h than at 24 h. In podoplanin-deficient keratinocytes, ppMLC2 staining appeared more prominent along stress fibres before stimulation and remained predominantly stress-fibre-associated at both time points, with little evident redistribution towards intercellular contacts (Figure11C). These differences were accompanied by defective reorganisation of the actin cytoskeleton. In control keratinocytes, Ca^2+^ stimulation promoted the progressive assembly of a robust cortical F-actin belt, which became increasingly evident at 24 and 48 h. Although cortical F-actin was also detected in podoplanin-deficient cells, it was less regularly organised and did not form a comparably robust and continuous cortical structure. Instead, mutant monolayers retained a more irregular and disorganised actin architecture (Figure11B-C).

**Figure 11. -.**
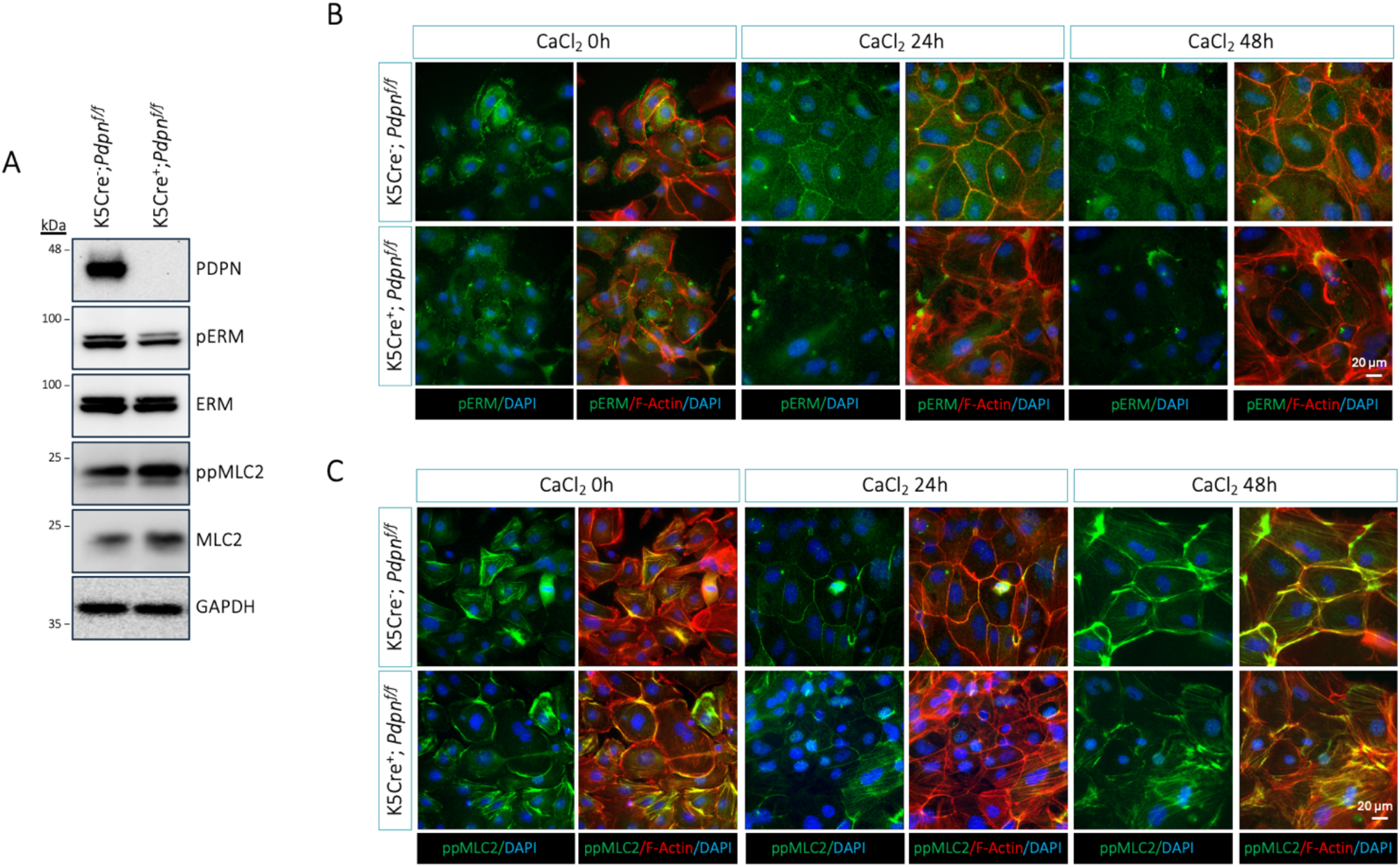
Ca^2+^-induced differentiation reveals altered ERM and MLC2 regulation in podoplanin-deficient keratinocytes. Primary keratinocytes isolated from K5Cre^-^;*Pdpn^f/f^* and K5Cre^+^;*Pdpn^f/f^* mice were induced to differentiate by supplementation with 1.2 mM CaCl₂. *(A)* Western blot analysis of podoplanin, phosphorylated ERM (pERM), total ERM, phosphorylated MLC2 (ppMLC2; Thr18/Ser19) and total MLC2 in keratinocytes before Ca^2+^ addition. GAPDH was used as a loading control. *(B)* Representative immunofluorescence images showing the distribution of pERM (green) and F-actin (red) before Ca^2+^ addition and after 24 and 48 h of differentiation. *(C)* Representative immunofluorescence images showing the distribution of ppMLC2 (green) and F-actin (red) at the indicated time points. Nuclei were counterstained with DAPI (blue). Podoplanin-deficient keratinocytes showed reduced pERM levels and increased basal ppMLC2, accompanied by altered redistribution of these proteins during Ca^2+^-induced differentiation. Western blots and images are representative of more than three independent keratinocyte isolations. Representative images from at least 4 independent experiments are shown. Scale bars, 20 µm.

Collectively, these findings show that podoplanin-deficient keratinocytes largely retain the ability to recruit junctional proteins to cell-cell contacts during Ca^2+^-induced differentiation, despite altered expression dynamics. However, these contacts display irregular boundaries and disorganised cortical F-actin, accompanied by reduced ERM phosphorylation and impaired junctional enrichment of pERM and ppMLC2. These results support a role for podoplanin in organising the junctional cortex during keratinocyte differentiation.

### Epidermal podoplanin loss alters the kinetics of TPA-induced skin remodelling

In homeostatic adult skin, podoplanin is absent from the interfollicular epidermis, with epithelial expression restricted to the outer root sheath (ORS) of anagen hair follicles (Yoon et al., 2019) (Figure 1B). However, it is induced in basal interfollicular keratinocytes and dermal fibroblasts in response to inflammatory or regenerative stimuli, including topical TPA treatment (Gandarillas et al., 1997; Montero-Montero et al., 2020). Because constitutive podoplanin KO mice die perinatally, we used the conditional K5CrePDPN model to investigate the specific contribution of keratinocyte-expressed podoplanin to the adult cutaneous response to TPA. Thus, an acute proliferative and inflammatory response was induced by applying 5 µg TPA topically to the dorsal skin in three consecutive doses. Mice were euthanised at 24, 48, 72 and 120 h after the third and final TPA application to monitor the temporal evolution of the cutaneous response from its early post-treatment phase towards resolution (Figure 12A).

**Figure 12. -.**
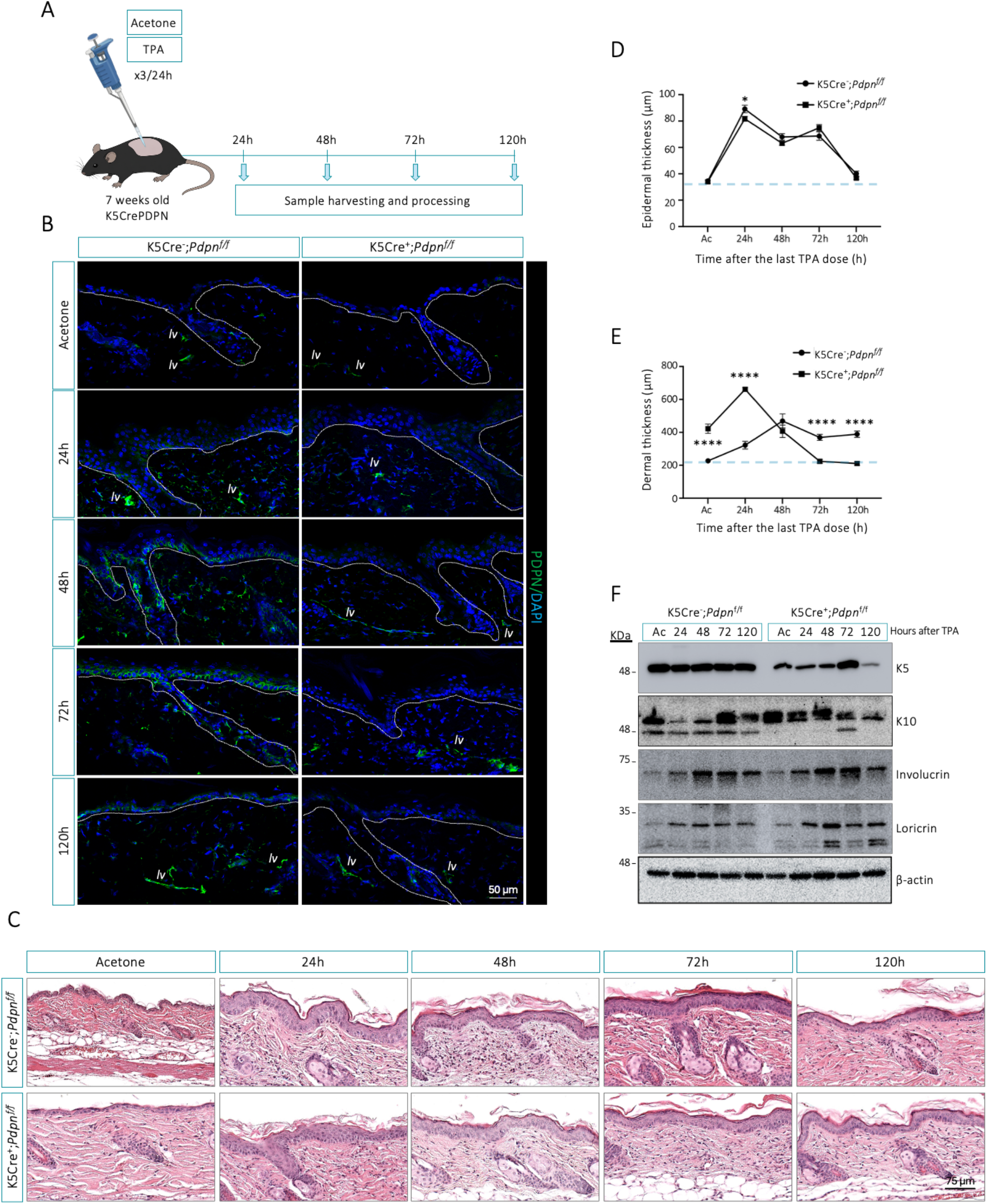
Altered kinetics of epidermal and dermal remodelling following acute TPA treatment in K5CrePDPN mice. *(A)* Schematic representation of the experimental protocol. Dorsal skin from 7-9-week-old male K5Cre^-^;*Pdpn^f/f^* and K5Cre^+^;*Pdpn^f/f^* mice was treated topically with acetone or TPA once daily for three consecutive days. Samples were collected at the indicated times after the final application. *(B)* Representative immunofluorescence images showing podoplanin expression (green) in dorsal skin from acetone- and TPA-treated mice. Nuclei were counterstained with DAPI (blue). White lines delimit the epidermis and hair follicles, and *lv* indicates dermal lymphatic vessels. TPA induced podoplanin expression in the control epidermis and dermal fibroblasts, whereas both responses were altered in K5Cre^+^;*Pdpn^f/f^* skin. *(C)* Representative H&E-stained dorsal skin sections collected after acetone treatment or at the indicated times following the final TPA application. *(D-E)* Time course of epidermal *(D)* and dermal *(E)* thickness following acute TPA treatment. Horizontal dashed lines indicate the mean basal value in acetone-treated K5Cre^-^;*Pdpn^f/f^* controls. *(F)* Representative western blot analysis of K5, K10 and involucrin in whole-skin lysates collected at the indicated times. β-Actin was used as a loading control. Data are presented as mean ± SEM (*n* ≥ 4 mice per group). Statistical comparisons between genotypes at each time point were performed using two-way ANOVA followed by Šídák’s multiple-comparisons test. *P* < 0.05 and \*\*\**P* < 0.0001. Scale bars, 50 µm in *(B)* and 75 µm in *(C)*.

Podoplanin expression was monitored throughout this time course. Consistent with previous reports, in control skin podoplanin was strongly induced in basal interfollicular keratinocytes following TPA treatment, reaching its highest levels between 48 and 72 h after the final application, and declining by 120 h (Figure 12B). Dermal fibroblasts also exhibited inducible podoplanin expression, with the strongest staining detected earlier, between 24-48 h. As expected, no epidermal podoplanin induction was detected in K5Cre^+^;*Pdpn^f/f^* mice at any time point. Unexpectedly, however, podoplanin induction in dermal fibroblasts was markedly attenuated in conditional KO skin, despite genetic deletion being restricted to keratinocytes. Histological analysis revealed a marked but reversible epidermal hyperplasia in both genotypes. Epidermal thickness reached its maximum 24 h after the final TPA application and progressively returned towards basal levels by 120 h (Figure 12C). Despite this similar overall temporal profile, epidermal thickness at 24 h was slightly but significantly lower in K5Cre^+^;*Pdpn^f/f^* mice than in their control counterparts, whereas no significant differences between genotypes were detected at later time points (Figure 12D). The dermal compartment displayed a distinct temporal response (Figure 12E). Under basal conditions, dermal thickness was already increased in podoplanin-deficient mice. Following TPA treatment, control skin showed a progressive increase in dermal thickness and overall cellularity, reaching its highest levels at 48 h and remaining elevated thereafter. In K5CrePDPN skin, the dermal response occurred earlier, reaching its maximum at 24 h before progressively declining to below basal levels by 120 h. Thus, keratinocyte-specific podoplanin loss altered both the timing and persistence of the dermal response to TPA.

We next analysed basal (K5), suprabasal (K10), intermediate (involucrin) and terminal (loricrin) differentiation markers by Western blotting and immunofluorescence at the indicated time points (Figure 12F and S3). In vehicle-treated skin, K5Cre^+^;*Pdpn^f/f^* epidermis showed reduced K5 expression, together with increased K10 and slightly elevated loricrin levels, indicating a pre-existing shift in the balance between basal and differentiating keratinocytes. Following TPA treatment, control epidermis displayed an ordered temporal response. K5 expression remained relatively stable throughout the time course, whereas K10 underwent a transient reduction at 24-48 h before increasing at later stages, consistent with the established repression of K1/K10 expression following TPA-mediated PKC activation (Dlugosz and Yuspa, 1993), before increasing at later time points. Involucrin expression increased progressively, while loricrin reached its highest level at 48 h and subsequently declined at 72-120 h. In podoplanin-deficient epidermis, these dynamics were disrupted. K5 remained comparatively reduced throughout the time course, whereas K10, which was already elevated under basal conditions, failed to undergo the early TPA-induced reduction observed in controls (Figure 12F and S3). Involucrin showed a stronger induction, while loricrin followed a biphasic temporal profile, characterised by an early peak at 24 h, a marked reduction at 48 h and renewed accumulation at 72-120 h, reaching levels above those observed in control skin (Figure 12F and S3).

Overall, these findings indicate that podoplanin deficiency in basal keratinocytes disrupts epidermal homeostasis and alters the kinetics of the inflammatory response. Rather than simply enhancing differentiation, podoplanin loss perturbs the coordinated progression of the epidermal differentiation programme under both basal and inflammatory conditions, consistent with the altered dynamics observed during skin morphogenesis *in vivo* and in Ca^2+^-induced differentiation *in vitro*. In parallel, the attenuated induction of podoplanin in dermal fibroblasts and the altered kinetics of dermal remodelling support the existence of non-cell-autonomous consequences of epidermal podoplanin deficiency. Together, these results support a role for podoplanin as an important modulator of epidermal remodelling and epidermal-dermal communication under conditions of tissue challenge.

### Slow-cycling epidermal cell populations show altered homeostasis and TPA responsiveness in podoplanin-deficient skin

Adult epidermal homeostasis and regeneration are sustained by distinct stem and progenitor populations located in the basal layer of the interfollicular epidermis (IFE) and in defined hair follicle (HF) niches. During homeostasis, these populations largely maintain their respective compartments, whereas injury or inflammatory stimulation can induce their exit from quiescence and proliferative expansion to support tissue remodelling and restoration of epithelial integrity (Gonzales and Fuchs, 2017; Sun et al., 2023). Given that podoplanin is re-expressed by basal keratinocytes during tissue remodelling in adult skin and its deficiency disrupts keratinocyte cytoskeletal organisation and division orientation during epidermal morphogenesis, we hypothesised that podoplanin might also influence the proliferative responsiveness of epidermal stem and progenitor populations during adult epidermal remodelling. To investigate this possibility, we combined the TPA-induced inflammatory model described above, which elicits a robust hyperproliferative and remodelling response, with a BrdU label-retaining cell (LRC) assay in K5CrePDPN mice. This experimental approach has previously been used to examine the activation of epidermal stem cell populations *in vivo* (Cotsarelis et al., 1990; Sotiropoulou et al., 2008). Thus, mice received BrdU during the early postnatal period (P6-P9) to label proliferating epidermal cells, followed by a chase period during which the label was progressively diluted in actively cycling cells but retained in slow-cycling populations (Figure 13A). After completion of the chase, adult mice were subjected to the TPA treatment protocol described above, and dorsal skin was collected 24 h after the final application. Long-term BrdU retention was used to identify slow-cycling epidermal populations enriched in cells with stem and progenitor properties. Accordingly, a TPA-induced reduction in label retention was considered consistent with their proliferative activation and consequent dilution of the BrdU label through successive cell divisions.

**Figure 13. -.**
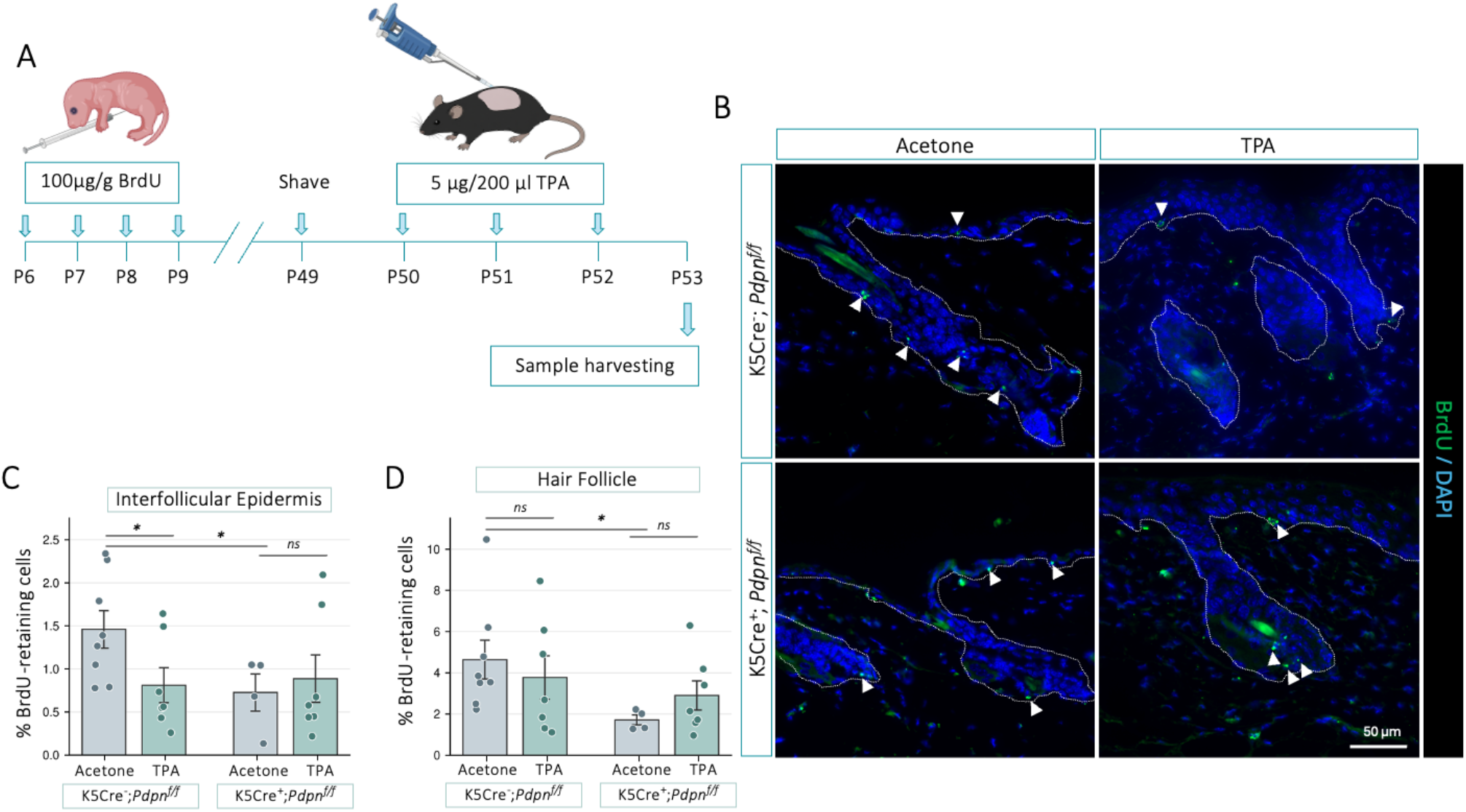
Epidermal podoplanin modulates label-retaining cell mobilisation in response to TPA. *(A)* Schematic representation of the BrdU pulse-chase protocol. Mice received daily BrdU injections (100 µg/g body weight) from P6 to P9, followed by a 41-day chase ending at P50. Dorsal skin was shaved at P49, and acetone or TPA (5 µg in 200 µl acetone) was applied once daily from P50 to P52. Skin samples were collected at P53, 24 h after the final application. *(B)* Representative immunofluorescence images of BrdU-retaining cells in 10µm dorsal skin sections from acetone- and TPA-treated K5Cre^-^;*Pdpn^f/f^* and K5Cre^+^;*Pdpn^f/f^* mice. BrdU is shown in green and DAPI in blue. Dashed white lines delimit the interfollicular epidermis (IFE) and hair follicles (HF), and arrowheads indicate BrdU-positive nuclei. *(C-D)* Percentage of BrdU-retaining cells among total DAPI-positive nuclei in the IFE *(C)* and HF *(D)*. Percentages obtained from individual fields were averaged to generate one value per mouse for each compartment. Data are presented as mean ± SEM, with each point representing an individual mouse (*n* = 4-8 mice per group). Prespecified comparisons were performed using two-tailed unpaired Welch’s *t*-tests. Unadjusted *P* values are shown. ns, not significant; \**P* < 0.05. Scale bar, 50 µm.

As shown in Figure 13B-D, TPA treatment significantly reduced the proportion of BrdU-retaining cells in the IFE of control mice, whereas HFs showed only a slight, non-significant decrease. The reduction in interfollicular label retention is consistent with cell-cycle entry and subsequent dilution of BrdU, suggesting activation of slow-cycling epidermal cells in response to TPA. However, no corresponding reduction was observed in the IFE or HF of podoplanin-deficient mice. Notably, vehicle-treated K5CrePDPN mice already exhibited fewer BrdU-retaining cells in both compartments than controls, indicating that differences in these populations were also present before inflammatory stimulation. Together, these findings suggest that podoplanin loss alters both the basal abundance or label-retention dynamics of slow-cycling epidermal populations and their response to TPA. These results extend the role of podoplanin beyond epidermal morphogenesis and support its contribution to the regulation of epidermal cell dynamics during adult tissue remodelling.

## MATERIAL AND METHODS

### Mouse strains and breeding

Podoplanin-floxed (*Pdpn^f/f^*) mice have been previously described (Lowe et al., 2015), and K5Cre transgenic mice were kindly provided by Dr. Ángel Ramírez (Ramirez et al., 2004). Both mouse strains were maintained on a C57BL/6 background. *Pdpn^f/f^* mice were crossed with K5Cre mice to generate mice lacking podoplanin in basal keratinocytes. K5Cre-positive breeders (K5Cre^+^) were maintained as hemizygotes throughout colony breeding.

Two experimental models were generated depending on the breeding strategy. Paternal transmission of K5Cre, through crosses between K5Cre^+^;*Pdpn^f/f^* males and K5Cre^−^; *Pdpn^f/f^* females, generated conditional epidermal KO mice (K5CrePDPN; K5Cre^+^;*Pdpn^f/f^*). Littermate K5Cre^−^;*Pdpn^f/f^* mice were used as controls. To generate the systemic constitutive KO model, K5Cre^−^;*Pdpn^w/f^*males were crossed with K5Cre^+^;*Pdpn^w/f^* females. Maternal K5Cre activity induced early and widespread recombination of the floxed *Pdpn* alleles, including in offspring that did not inherit the K5Cre transgene, as reported previously (Ramirez et al., 2004). Cre-negative offspring carrying two recombined alleles (K5Cre^−^;*Pdpn^Δ/Δ^*) were selected as systemic KO mice, whereas K5Cre^−^;*Pdpn^w/w^*littermates were used as controls. The presence of the recombined *Pdpn^Δ^* allele was confirmed by PCR in all KO animals as described below (Figures 2-3 and Table S1). Adult experiments were restricted to male mice to standardise the experimental cohorts and limit potential variation associated with sex-related differences in hormonal status and hair follicle cycling. Embryos and neonates were analysed irrespective of sex. Embryos were collected between embryonic day (E)12.5 and E18.5. Postnatal analyses were performed in neonates at postnatal day (P)0 and P5 and in adult mice aged 7-9 weeks, corresponding to the telogen phase of the hair cycle.

Genotyping was performed by PCR using genomic DNA isolated from ear punch biopsies in adult mice and from tail tissue in embryos and neonates. The K5Cre transgene was detected using a specific oligonucleotide pair. A three-primer PCR strategy was used to discriminate among the wild-type (*Pdpn^w^*), floxed (*Pdpn^f^*) and recombined (*Pdpn^Δ^*) alleles (Figures 2-3 and Table S1).

Mice were housed in the Animal Facility of Universidad Francisco de Vitoria (ES281150000013) under controlled conditions (12 h light/dark cycle, ∼22 °C) with food and water provided ad libitum. Adult experimental mice and pregnant females were euthanised by CO₂ inhalation followed by cervical dislocation. Neonatal mice and embryos were euthanised by decapitation. All animal procedures were approved by the institutional ethics committee and authorised by the Comunidad de Madrid (PROEX 068.6/21), in accordance with European Directive 2010/63/EU and Spanish legislation (RD 53/2013) governing the use of animals in scientific research purposes.

### In vivo epidermal barrier assay

Epidermal permeability barrier formation during embryonic development was assessed using a toluidine blue dye-exclusion assay, as previously described (Schmitz et al., 2015). Briefly, mouse embryos were collected at embryonic stages E16.5-E18.5 and rinsed in cold phosphate-buffered saline (PBS). Embryos were sequentially dehydrated and rehydrated through a graded methanol series (25%, 50%, 75%, 100%, 75%, 50% and 25%; 1 min each), followed by rinsing in PBS. Samples were then incubated in 0.1% toluidine blue solution for 10 min at room temperature and extensively washed in PBS to remove excess dye. An intact epidermal barrier excluded the dye, whereas blue staining indicated incomplete or defective barrier formation. Images were acquired using a stereomicroscope (Leica EZ4) in lateral, dorsal and ventral orientations. The percentage of the visible embryo area lacking dye penetration was quantified separately for each orientation using Fiji/ImageJ software. A total of 56 embryos from 10 litters were analysed for the constitutive model and 52 embryos from seven litters for the conditional model across developmental stages E16.5-E18.5.

### Histological analysis

Embryos and skin samples were collected at the indicated developmental stages and fixed in 10% neutral-buffered formalin at 4 °C for 24-48 h. Embryos collected at E12.5-E14.5 were fixed for 24 h, whereas embryos collected from E16.5 onwards underwent a ventral incision to facilitate fixative penetration and were fixed for 48 h. Samples were processed without decalcification, dehydrated through graded ethanol solutions, cleared in xylene and embedded in paraffin. Paraffin-embedded samples were sectioned at 3-5 µm, typically 4 µm, using a microtome and mounted on glass slides. For general histological evaluation, sections were stained with haematoxylin and eosin (H&E) following standard procedures. Periodic acid-Schiff (PAS) staining combined with Alcian blue at pH 2.5 was performed on additional sections to evaluate basement membrane integrity at the dermal-epidermal junction. Whole-slide images were acquired at 20× magnification using an Aperio Versa slide scanner (Leica Biosystems) at the Histology Service of the Instituto de Medicina Molecular Aplicada (IMMA) Nemesio Díez, Universidad CEU San Pablo. Digital slides were examined using Aperio ImageScope software.

### Morphometric and proliferation analyses

Epidermal and dermal thicknesses were quantified in digital whole-slide images of H&E-stained dorsal skin sections from embryos, neonatal mice and adult mice using Aperio ImageScope software (Leica Biosystems). Interfollicular epidermal (IFE) thickness was determined by drawing lines perpendicular to the basement membrane, extending from the dermal-epidermal junction to the outer surface of the stratum corneum. Dermal thickness was measured using perpendicular lines extending from the basement membrane to the dermal-hypodermal junction. Approximately 60-80 measurements were performed per animal, distributed across approximately 3,000 µm of tissue. Measurements were averaged to generate a single epidermal or dermal thickness value for each animal, which was used as the experimental unit for statistical analysis.

Cell proliferation was assessed by Ki67 immunofluorescence staining. The proliferative index was calculated as the percentage of Ki67-positive nuclei relative to the total number of DAPI-positive nuclei within the IFE. Multiple fields were quantified per section using Fiji/ImageJ, and the values obtained were averaged to generate a single value for each animal for subsequent statistical analysis.

### Immunodetection Immunofluorescence staining

For immunofluorescence staining of cultured cells, cells were fixed in 3.7% formaldehyde for 15 min and permeabilised with 0.05% Triton X-100 for 10 min. Primary and secondary antibodies were diluted in PBS containing 0.1% bovine serum albumin (BSA), which also served as the blocking solution. Cells were incubated with the indicated primary antibodies overnight at 4°C (Table S2). After washing with PBS, cells were incubated with the corresponding species-specific fluorescent secondary antibodies for 1 h at room temperature and counterstained with DAPI (Sigma-Aldrich D9542). Coverslips were mounted using ProLong Gold Antifade Mountant (Invitrogen). Fluorescence images were acquired using a Leica THUNDER widefield microscope (Leica Microsystems) equipped with HC PL APO 40×/1.30 Oil and HC PL APO 63×/1.40–0.60 Oil objectives.

For immunofluorescence staining of tissue sections, paraffin sections were deparaffinised and rehydrated before heat-induced epitope retrieval in EDTA buffer (pH 8.0) using a pressure cooker. Sections were blocked in PBS containing 1% BSA and 2% fetal bovine serum (FBS) and incubated with the indicated primary antibodies (Table S2) overnight at 4 °C. After washing with PBS, sections were incubated with the corresponding fluorescent secondary antibodies for 1 h at room temperature and counterstained with DAPI. Slides were mounted using ProLong Gold Antifade Mountant (Invitrogen). Images were acquired using either a Leica THUNDER microscope equipped with an HC PL APO 40×/1.30 Oil or a Leica STELLARIS confocal microscope equipped with an HC PL APO CS2 20x/0.75. Images shown in Figures 12 and S3 were acquired using a Zeiss LSM 710 spectral confocal microscope equipped with a Plan-Apochromat 25×/0.8 Imm Korr DIC objective.

### Western blotting

Cells or tissue samples were lysed in 2% SDS buffer containing 50 mM Tris-HCl pH 7.4, 100 mM NaCl, 5 mM MgCl2, 5 mM CaCl2 and 2% SDS, and supplemented with pretease inhibitor cocktail set I (Calbiochem) and phosphatase inhibitor complex PhosSTOP from Roche. Protein concentrations were determined using the BCA assay (Pierce, Rockford, IL, USA) for cell lysates and the Bradford assay (Bio-Rad) for tissue lysates.

Equal amounts of protein (10-30 µg) were separated on 8%, 10% or 12.5% SDS-polyacrylamide gels, depending on the molecular weight of the protein of interest, and transferred to Immobilon-P PVDF membranes (Millipore). Membranes were blocked in Tris-buffered saline containing 0,1% Tween 20 (TBS-T) supplemented with 5% non-fat dry milk for total-protein detection or 3% BSA for phosphoprotein detection. Membranes were incubated with the indicated primary antibodies overnight at 4°C (Table S2), washed with TBS-T and incubated with the corresponding horseradish peroxidase-conjugated secondary antibodies for 1 h at room temperature. After further washing with TBS-T, immunoreactive bands were detected using an ECL chemiluminescent substrate (Pierce).

### Antibodies

Details of all primary and secondary antibodies used in this study, including supplier, catalogue number and working dilution, are provided in Supplementary Table S2.

### Mitotic division angle analysis

Mitotic division orientation was analysed in basal epidermal cells from embryonic skin sections. For the constitutive KO model, embryos were analysed at E12.5, E16.5 and E18.5, whereas the conditional KO model was analysed at E12.5, E17.5 and E18.5. Paraffin sections (10 μm) were immunostained for phospho-histone H3 (pHH3), β-catenin (Table S2) and counterstained with DAPI (Sigma-Aldrich D9542). Basal mitotic cells were identified by their location within the basal epidermal layer, with β-catenin staining used to delineate cell boundaries and the basal-dermal interface. Only pHH3-positive basal mitoses containing two clearly separated DAPI-positive chromatin masses and a readily identifiable division axis were included. The division axis was defined by a line connecting the centres of the two chromatin masses. The angle between the division axis and the local basement membrane plane was measured in Fiji/ImageJ. Divisions were classified as planar/symmetric (0° to <30°), oblique (30-60°), or perpendicular/asymmetric (>60° to 90°). Three embryos were analysed per genotype and developmental stage, corresponding to 37-81 mitoses per genotype and stage and 685 mitoses overall. The division angles from each genotype/stage were pooled and grouped into six 15° intervals between 0° and 90° and represented as polar histograms, in which the radial height of each bar indicates the fraction of all divisions within the corresponding angular interval. Bars were colour-coded according to the three predefined orientation categories, and the total number of mitoses included in each condition is indicated as *n*. For quantitative summaries, the percentages of planar, oblique and perpendicular divisions were first calculated independently for each embryo and subsequently expressed as the mean ± SEM of three embryos. Fluorescence images were acquired using a Leica THUNDER widefield microscope (Leica Microsystems) and an HC PL APO 63×/1.40–0.60 OIL objective.

### Acute topical TPA treatment

To induce acute epidermal remodelling, 7-9-week-old male mice, corresponding to the telogen phase of the hair cycle, were used. Dorsal hair was removed two days before treatment to minimise skin irritation caused by shaving. 12-O-Tetradecanoylphorbol-13-acetate (TPA; Sigma-Aldrich, P8139) was applied topically to the shaved dorsal skin at a dose of 5 μg per mouse in 200 μL of acetone. Treatment was administered once daily for three consecutive days. Vehicle-treated animals of each genotype received an equivalent volume of acetone alone. Mice were euthanised, and dorsal skin samples were collected 24, 48, 72 or 120 h after the final application (Figure 12A).

### BrdU pulse-chase label-retaining cell assay

Label-retaining cells (LRCs) were identified using a BrdU pulse-chase assay. Neonatal mice received intraperitoneal injections of BrdU (100 μg/g body weight, 30 μl final volume per pup) at postnatal days P6-P9. After the labelling period, mice were subjected to a 41-day chase, ending at postnatal day 50 (P50), during which BrdU was progressively diluted in proliferating cells but retained in slow-cycling cells. Following the chase period, mice were treated with topical TPA (three consecutive daily applications) as described above. Skin samples were collected 24 h after the final TPA application (Figure 13A).

Paraffin-embedded skin sections were processed for immunofluorescence detection of BrdU and keratin 5 (K5), with DAPI (Sigma-Aldrich D9542) used as a nuclear counterstain. K5 staining was used to identify the basal keratinocyte compartment of the interfollicular epidermis (IFE) and the epithelial regions of the hair follicles (HF), but the K5 channel is not shown to facilitate visualisation of the BrdU signal. LRCs were defined as BrdU-positive nuclei located within these K5-positive compartments. LRCs were quantified using a custom macro developed in Fiji/ImageJ. IFE and HF regions were manually delineated and analysed separately. DAPI-positive nuclei and BrdU-positive nuclei were identified by thresholding and automated image segmentation. The percentage of BrdU-positive nuclei was calculated for each field, and field values were averaged to obtain a single value per mouse for the IFE and HF. Data were plotted using GraphPad Prism as individual mouse values together with the mean ± SEM, with acetone- and TPA-treated animals grouped by genotype.

### Isolation and culture of primary keratinocytes

Primary mouse keratinocytes were isolated from the dorsal skin of postnatal day 3-4 pups. Skin samples were dissected and incubated overnight at 4 °C in Dispase II solution (2.5 U/mL; Roche, 04942078001). The following day, samples were incubated at 37 °C for 5-10 min, and the epidermis was carefully separated from the dermis. Epidermal sheets were incubated in 0.25% trypsin-EDTA for 5-10 min at 37 °C. Trypsin activity was neutralised with medium containing 5% FBS. The resulting cell suspension was passed through a 70 μm cell strainer, centrifuged and washed with PBS. Cell viability was assessed by trypan blue exclusion before plating. Keratinocytes were seeded onto culture plates coated with 10 μg/mL Corning® UltraPure™ Laminin (Corning, 354239) and maintained in CnT-Prime Epithelial Proliferation Medium (CnT-PR; CELLnTEC). Cells were cultured at 37 °C in a humidified atmosphere containing 5% CO₂.

### Ca^2+^-induced keratinocyte differentiation

Keratinocyte differentiation was induced by replacing the proliferation medium with CnT-Prime Epithelial 2D Differentiation Medium (CnT-PR-D; CELLnTEC), adjusted to a final CaCl₂ concentration of 1.2 mM. Cells were maintained under differentiation conditions for 24, 48 and/or 120 h, depending on the experiment.

## Statistical analysis

Statistical analyses were performed using GraphPad Prism 10.0 (GraphPad Software). Data normality was assessed using the Shapiro-Wilk test. Comparisons between two independent groups were performed using two-tailed unpaired Student’s t-tests for normally distributed data. In the LRC experiments, two-tailed unpaired Welch’s t-tests were used to account for potentially unequal variances between groups. Experiments involving two independent variables, such as genotype and developmental stage or time after TPA treatment, were analysed using two-way ANOVA followed by Šídák’s multiple comparisons test. Measurements obtained from multiple fields, cells or tissue regions were averaged to obtain a single value for each biological replicate before statistical analysis.

Data are presented as mean ± standard error of the mean (SEM), with individual biological replicates shown where indicated. Differences were considered statistically significant at p < 0.05. Statistical significance is denoted as follows: *p < 0.05, **p < 0.01, ***p < 0.001 and ****p < 0.0001.

## DISCUSSION

Here, by comparing global podoplanin deletion with K5Cre-mediated deletion in basal keratinocytes, we define a previously unrecognised function for podoplanin during epidermal morphogenesis, showing that it supports keratinocyte structural organisation and contributes to epidermal architecture, differentiation and barrier acquisition. In adult skin, podoplanin contributes to the response to the pro-inflammatory agent TPA, influencing both label-retaining cell dynamics and epidermal and dermal remodelling. Together, these findings establish podoplanin as a regulator of skin organisation during periods of active tissue reorganisation, including development and inflammatory remodelling, rather than as a constitutive requirement for basal maintenance of the adult interfollicular epidermis. Podoplanin is a mucin-like transmembrane glycoprotein best known for its roles in lymphatic vascular development, and CLEC-2-dependent communication with platelets and immune cells (Quintanilla et al., 2019). In the skin, epidermal podoplanin expression has primarily been associated with contexts of active tissue remodelling, including inflammation, wound repair, psoriasis-like hyperproliferation and squamous cell carcinoma (SCC) progression (Gandarillas et al., 1997; Honma et al., 2012; Martín-Villar et al., 2005). Although functional studies, largely performed in cultured keratinocytes and SCC cells, have shown that podoplanin can modulate cell adhesion, migration and plasticity (Martín-Villar et al., 2006, 2005; Shibuya et al., 2019), its physiological contribution to skin development and homeostasis remains poorly understood. *In vivo* studies using conditional KO models have largely focused on adult skin, where podoplanin is barely expressed in the interfollicular epidermis under homeostatic conditions and its deletion in keratinocytes produces no overt epidermal phenotype (Baars et al., 2015; Sesartić et al., 2020; Yoon et al., 2019). Its role during epidermal morphogenesis has received little attention, while its contribution to inflammatory skin remodelling remains largely unexplored despite its induction in keratinocytes during inflammation.

To our knowledge, this study provides the first detailed characterisation of the spatiotemporal dynamics of podoplanin expression during epidermal morphogenesis. Podoplanin was detected in the early embryonic epidermis and became progressively enriched in basal epithelial compartments engaged in active growth and remodelling, a distribution reminiscent of activated epidermal stem and progenitor cell populations. During the early postnatal period, its expression declined in the interfollicular epidermis while remaining prominent in growing hair follicles, particularly within the outer root sheath (ORS). Both global and K5Cre-mediated podoplanin deletion caused keratinocyte vacuolisation, basal layer disorganisation, increased epidermal thickness and proliferation at late embryonic stages, and altered distribution of differentiation markers. This shared phenotype supports a direct contribution of keratinocyte-expressed podoplanin to epidermal morphogenesis, whereas the additional dermal oedema and vascular congestion in the global KO likely reflect its systemic nature (Mahtab et al., 2008; Schacht et al., 2003). Barrier acquisition was delayed at E16.5-E17.5, but dye exclusion was largely restored by E18.5 despite persistent histological and differentiation abnormalities. This recovery indicates that podoplanin is not indispensable for the eventual barrier acquisition but does not imply complete epidermal maturation. The accompanying structural and differentiation defects support a role for podoplanin in coordinating epidermal organisation with timely functional maturation. Previous studies using constitutive epithelial-specific deletion models have focused primarily on adult skin. K14Cre-mediated podoplanin deletion did not impair gross re-epithelialisation following acute full-thickness wounding (Baars et al., 2015), whereas K5Cre-mediated deletion produced no obvious abnormalities in the adult interfollicular epidermis under homeostatic conditions (Yoon et al., 2019). Our findings extend these observations by revealing developmental epidermal defects that become less pronounced at later stages. This attenuation partly reflects compensatory mechanisms during tissue maturation and may help explain why these abnormalities went unnoticed in studies focused on adult skin. We also found that follicular dynamics were affected: neonatal podoplanin-deficient skin exhibited delayed hair follicle differentiation, while earlier anagen entry and accelerated hair regrowth following depilation were previously reported in adult K5Cre-mediated deletion models (Yoon et al., 2019). Together, these findings support a role for podoplanin in the regulation of hair follicle development and cycling. Its declining expression in adult skin and the attenuation of the IFE phenotype with maturation further suggest that its contribution is particularly important while epidermal architecture is being established.

This role in developmental coordination also extends to epidermal differentiation. Podoplanin-deficient epidermis showed altered K5 expression, expanded and aberrant K10 expression, together with premature loricrin expression, while primary mutant keratinocytes underwent earlier morphological changes and displayed altered kinetics of differentiation-associated proteins following Ca^2+^ addition. Previous studies in human epidermis linked podoplanin expression to impaired granular layer and filaggrin formation (Honma et al., 2013), and subsequent work reported its downregulation during keratinocyte differentiation (Honma et al., 2019). Our findings show that this relationship cannot be explained simply by podoplanin acting as an inhibitor of differentiation: altered distribution and premature appearance of differentiation markers was accompanied by parakeratosis, cornified layer abnormalities and delayed barrier acquisition. Thus, altered differentiation marker expression did not translate into timely barrier acquisition. Instead, these observations suggest that podoplanin helps coordinate the transition from basal identity to terminal differentiation with the structural maturation required for barrier function.

Such coordination depends on the integration of basal cell shape, cortical tension, cell-cell and cell-matrix adhesion, division orientation and delamination during stratification (Damen et al., 2021; Miroshnikova et al., 2018; Simpson et al., 2011). Epidermal models lacking SRF or Plexin-B1/B2 display defects in cortical organisation, basal cell architecture and division orientation that overlap with those observed in podoplanin-deficient epidermis (Jiang et al., 2021; Luxenburg et al., 2011; Verdoni et al., 2010; Yin et al., 2023). These similarities support the view that podoplanin contributes to epidermal morphogenesis by regulating actin cortex organisation and basal cell architecture. In both podoplanin-deficient models, the developmental shift towards perpendicular basal divisions was attenuated, with more oblique divisions in the global KO and persistent planar orientations following K5Cre-mediated deletion. Although division angles alone do not establish daughter cell fate, their distribution contributes to epidermal growth and stratification (Lechler and Fuchs, 2005; Williams et al., 2011). Ca^2+^-induced differentiation further revealed altered regulation of components of adherens junctions, tight junctions and desmosomes. Podoplanin-deficient keratinocytes exhibited higher basal levels of E-cadherin and desmoglein-1, together with altered expression dynamics of E-cadherin and claudin-1 during differentiation. Recruitment of E-cadherin, β-catenin, ZO-1 and claudin-1 to cell-cell contacts was largely preserved, but ZO-1 and claudin-1 showed increased cytoplasmic accumulation at baseline and 24 h after Ca^2+^ addition. Thus, the junctional phenotype involved changes in both protein abundance and subcellular distribution, accompanied by irregular cell boundaries and disorganised cortical F-actin. These findings suggest that podoplanin contributes to the coordination of junctional protein regulation with cortical reorganisation during keratinocyte differentiation. Previous work from our group established that podoplanin interacts with ERM proteins and regulates their subcellular distribution, RhoA/C-ROCK-LIMK signalling and actin cytoskeletal remodelling in epithelial cells (Martín-Villar et al., 2015, 2006). A functional connection between podoplanin and actomyosin contractility has also been demonstrated in fibroblastic reticular cells (FRCs) (Astarita et al., 2015; Makris et al., 2024). Here, the altered organisation of the junctional cortex in podoplanin-deficient keratinocytes was accompanied by reduced pERM, increased basal MLC2/ppMLC2, and failure to concentrate both pERM and ppMLC2 at cell-cell contacts during Ca^2+^-induced differentiation. Thus, the irregular cell boundaries cannot be attributed to a simple loss of junctional proteins or a general decrease in actomyosin activity. Rather, the combined alterations in cortical F-actin, pERM and ppMLC2 indicate that podoplanin contributes to the spatial organisation of mechanical activity at the junctional cortex. To our knowledge, these findings provide the first evidence that podoplanin regulates the junctional enrichment of both pERM and ppMLC2 during keratinocyte differentiation, extending its established role in cytoskeletal remodelling to the organisation of the cortex at epithelial cell-cell contacts. This interpretation is consistent with the close interdependence between cell adhesion, cortical tension and barrier formation in stratified epithelia (Rübsam et al., 2017; Sumigray and Lechler, 2015; Tunggal et al., 2005).

The response to TPA in K5CrePDPN mice reinforces the idea that podoplanin becomes functionally important during active tissue reorganisation in the adult. Podoplanin/PA2.26 was initially characterised as a cell-surface antigen induced by TPA in epidermal keratinocytes and fibroblasts during chemical skin carcinogenesis (Gandarillas et al., 1997; Martín-Villar et al., 2010; Montero-Montero et al., 2020) and subsequently identified as a transcriptional target of Fos/AP-1 (Durchdewald et al., 2008) in the same context. Accordingly, in the present study, podoplanin was re-expressed in the control epidermis after acute TPA treatment, confirming that adult keratinocytes reactivate a developmental protein programme during inflammatory remodelling. Epidermal deletion modified not only the epidermal response to TPA but also the behaviour of the underlying dermis. The overall course of epidermal thickening was broadly preserved, although its early kinetics and the temporal expression of differentiation markers were altered. In contrast, dermal thickness displayed a markedly different temporal profile, and induction of podoplanin in dermal fibroblasts was reduced despite its preserved expression in lymphatic vessels. Because recombination was restricted to K5-expressing epithelial cells, these results reveal that epidermal podoplanin influences the response of neighbouring dermal cells. Podoplanin therefore participates in epidermal-dermal communication during cutaneous remodelling, extending its function beyond the cell-autonomous regulation of keratinocyte architecture. Although BrdU retention does not define a single molecularly distinct stem-cell population, the analysis of label-retaining cells (LRCs) provides an additional link between podoplanin and the response of long-lived, slow-cycling epidermal populations. Under acetone-treated conditions, K5CrePDPN skin contained fewer BrdU-retaining cells in both the interfollicular epidermis and hair follicles. TPA reduced LRC retention in the control interfollicular epidermis, consistent with activation or mobilisation of these cells, whereas this response was absent in the KO. In the hair follicle compartment, no significant reduction was detected under our experimental conditions, although controls showed a downward trend that was not apparent in K5CrePDPN mice. Quantification of the entire follicle may have masked redistribution between the bulge and other follicular regions. Together, these observations indicate that podoplanin influences LRC dynamics, affecting both their abundance after the chase period and their response to an acute proliferative stimulus.

Collectively, our findings identify a physiological role for keratinocyte-expressed podoplanin in coordinating epidermal organisation and functional maturation during development and adult tissue remodelling. At the molecular level, this role is associated with the regulation of ERM phosphorylation and its recruitment to cell-cell contacts together with ppMLC2, linking podoplanin to the cortical machinery that supports epithelial architecture. In adult skin, altered LRC dynamics following TPA extend its influence on the response of slow-cycling epidermal populations, while the accompanying dermal changes suggest that its effects may also involve epidermal-dermal communication. Together, these findings place podoplanin at the interface between keratinocyte organisation and the wider tissue response, providing a basis for investigating its contribution to inflammatory, reparative and neoplastic skin remodelling.

## ETHICS STATEMENT

All animal procedures were approved by the institutional ethics committee of Universidad Francisco de Vitoria and authorised by the Comunidad de Madrid (PROEX 068.6/21). Experiments were conducted in accordance with European Directive 2010/63/EU and Spanish Royal Decree 53/2013 on the protection of animals used for scientific purposes.

## DATA AVAILABILITY STATEMENT

All data supporting the findings of this study are included in the manuscript and supplementary materials or are available from the corresponding author upon request.

## ACKNOWLEDGEMENTS

This work was supported by grant PID2021-126816OB-I00 awarded to E.M.-V. by the Spanish Ministry of Science and Innovation (MCIN) and the State Research Agency (AEI/10.13039/501100011033), co-funded by the European Regional Development Fund (ERDF, European Union), and by research grants from Universidad Francisco de Vitoria (UFV2022-10, UFV2024-21 and UFV2026-34). R.J. was supported by an FPU predoctoral fellowship from the Spanish Ministry of Universities (FPU2022-03481). S.G. was supported by a predoctoral research contract funded by Universidad Francisco de Vitoria (IPF-UFV programme). B.G.-H. was supported by a research contract funded by the Comunidad de Madrid through its Programa de Jóvenes Investigadores (2025). We thank the Histology Facility at CNB/CSIC for the histological processing of biological samples. We also thank Drs. I. Antón and Y. Calle for helpful discussions of the results during manuscript preparation.

## AUTHOR CONTRIBUTIONS

R.J. performed carried out most of the experimental work, analysed and interpreted the data, contributed to figure preparation and critically reviewed the manuscript. S.T.-C. managed the mouse colonies, performed experiments and critically reviewed the manuscript. A.B. performed the histopathological assessment of the mouse phenotypes and critically reviewed the manuscript. S.G.-F., B.G.-B., M.G. and S.R. contributed to experimental work, including immunofluorescence assays. D.M. provided advice on confocal microscopy and acquisition of confocal images. S.B. and J.Mi. provided advice on intercellular junctions, contributed to data analysis and critically reviewed the manuscript. P.F., V.G.-G. and J.Mo. processed samples for histology. S.A.L., L.M.-M. and E.G. contributed to experimental work and critically reviewed the manuscript. S.P.W. provided the *Pdpn*-floxed mice and critically reviewed the manuscript. E.M.-V. conceived and designed the study, performed experiments, analysed and interpreted the data, prepared figures, supervised the project, secured funding and wrote the manuscript.

## DECLARATION OF GENERATIVE ARTIFICIAL INTELLIGENCE (AI) OR LARGE LANGUAGE MODELS (LLMs)

During the preparation of this manuscript, the authors used ChatGPT (OpenAI) to improve the English language, clarity and readability of the text. All AI-assisted content was carefully reviewed and edited by the authors, who take full responsibility for the final content of the manuscript.

## Supplementary Materials

**Figure S1. -.**
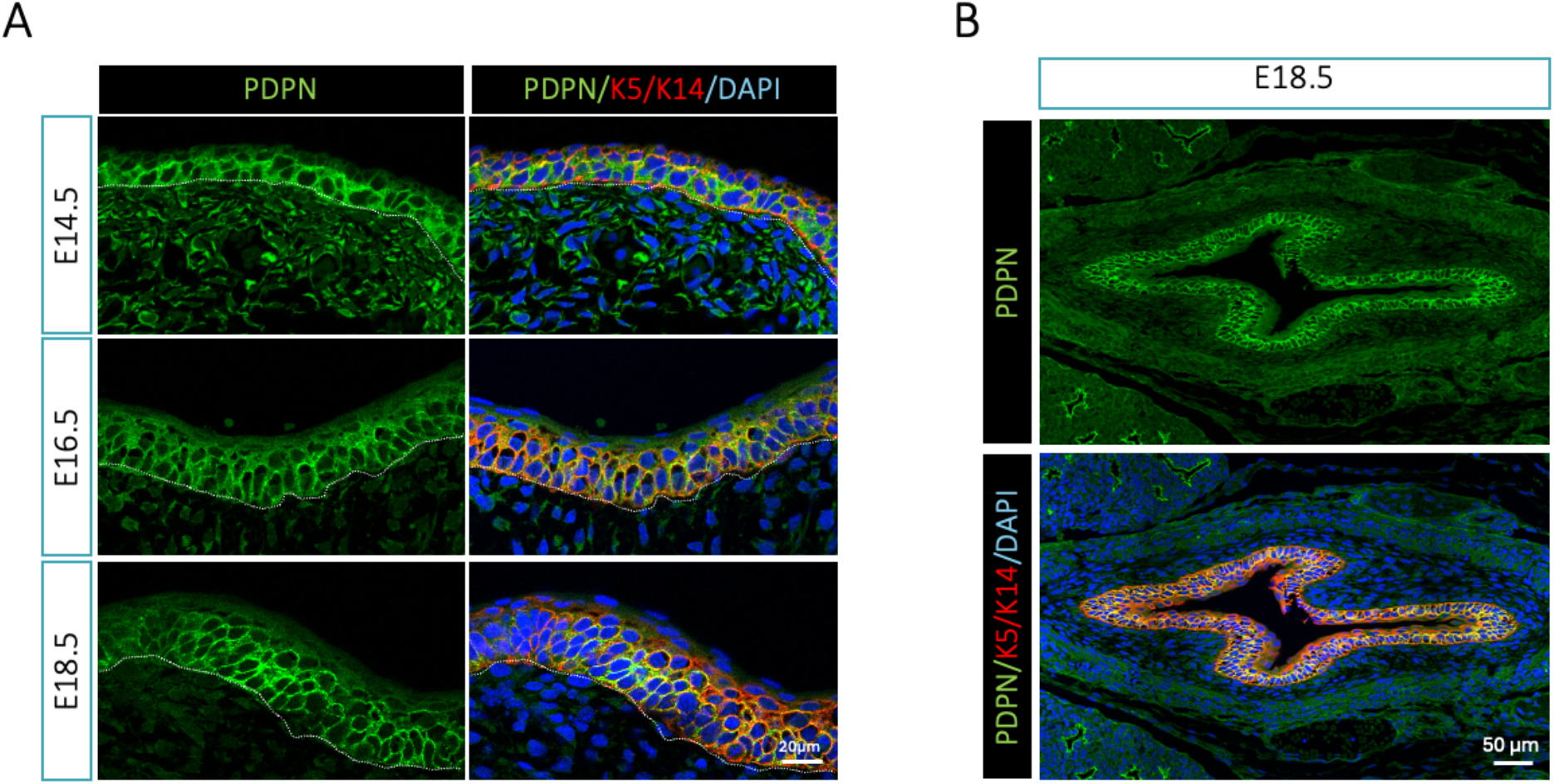
Developmental expression of podoplanin in the hard palate and oesophageal epithelium. *(A)* Representative confocal images showing podoplanin expression in the hard palate epithelium of wild-type C57BL/6 embryos at E14.5, E16.5 and E18.5. Sections were stained for podoplanin (green) and K5/K14 (red), with DAPI nuclear counterstaining (blue). The dashed white line indicates the dermo-epithelial boundary. *(B)* Representative transverse section of the oesophagus from a wild-type C57BL/6 embryo at E18.5 showing podoplanin expression (green) in the stratified epithelium. Representative images are shown for each developmental stage (n = 3-4 animals per stage). Scale bars, 20 µm in *(A)* and 50 µm in *(B)*.

**Figure S2. -.**
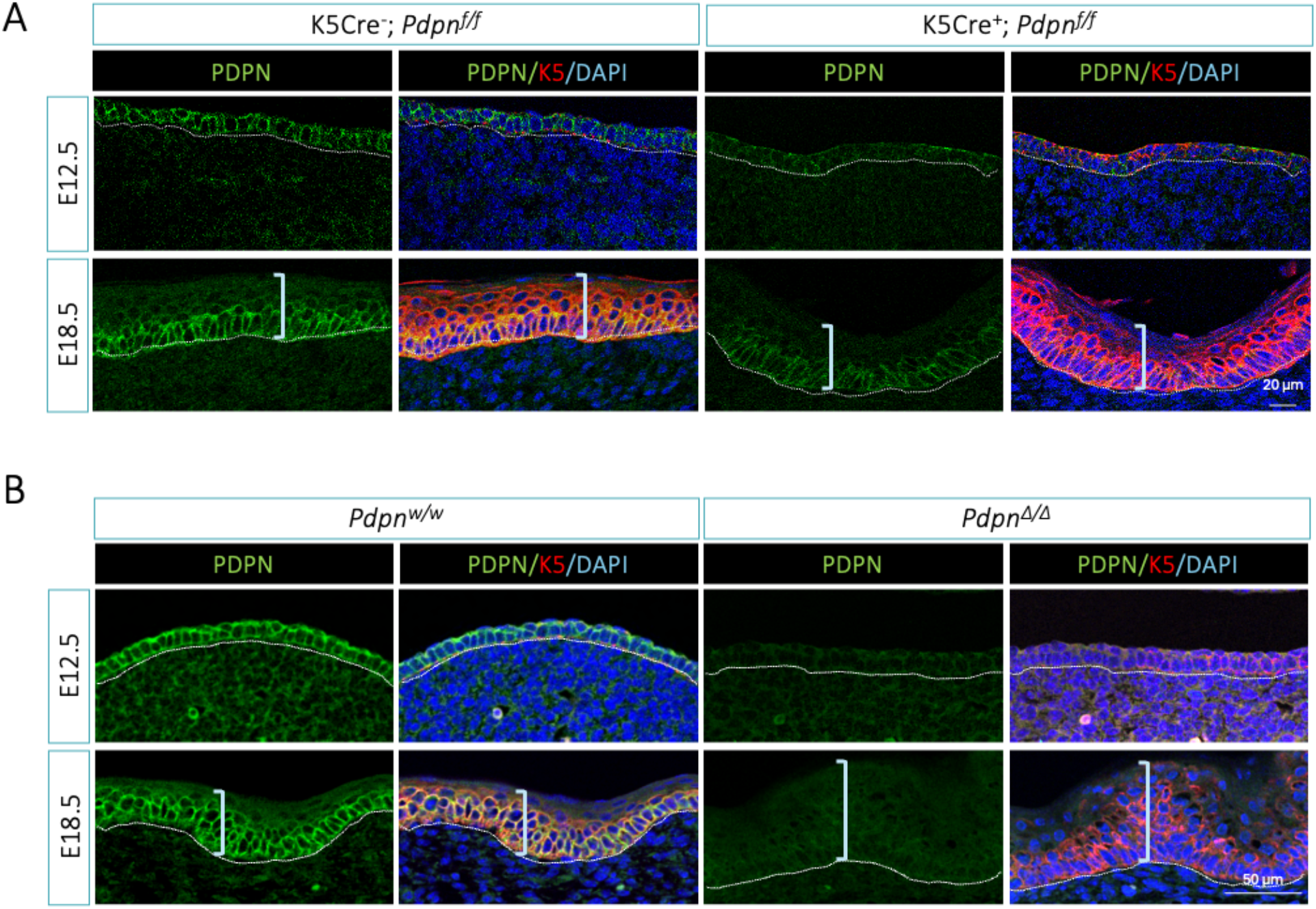
Validation of podoplanin deletion in the developing hard palate. *(A)* Representative confocal images showing podoplanin expression in the hard palate epithelium of K5Cre^-^;*Pdpn^f/f^* and K5Cre^+^;*Pdpn^f/f^* embryos at E12.5 and E18.5. *(B)* Podoplanin expression in the hard palate epithelium of *Pdpn^w/w^* and *Pdpn^Δ/Δ^* embryos at the indicated developmental stages. Sections were stained for podoplanin (green) and K5 (red), with DAPI nuclear counterstaining (blue). Dashed white lines indicate the dermo-epithelial boundary. The conditional model shows a progressive loss of epithelial podoplanin expression, whereas podoplanin is absent from the earliest stage analysed in the constitutive model. Representative images are shown for each developmental stage (n = 3-4 animals per stage). Brackets indicate oral epithelial thickness. Scale bars, 20 and 50 µm.

**Figure S3. -.**
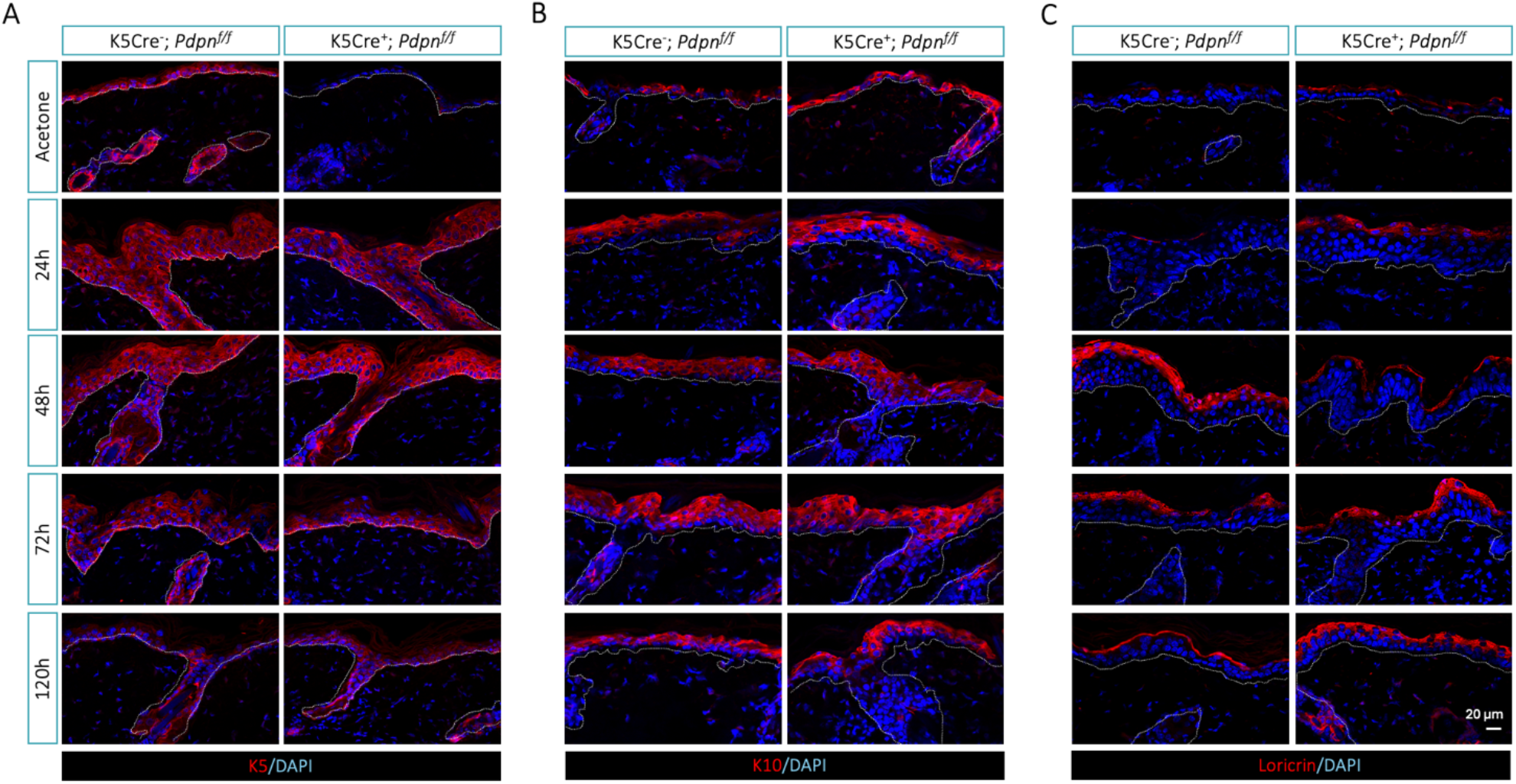
Spatiotemporal expression of epidermal differentiation markers following acute TPA treatment in K5CrePDPN skin. Representative confocal images of dorsal skin immunofluorescence from K5Cre^-^;*Pdpn^f/f^* and K5Cre^+^;*Pdpn^f/f^* mice treated with acetone or collected at 24, 48, 72 and 120 h after the final TPA application. Sections were stained for K5 (A), K10 (B) or loricrin (C), shown in red, with DAPI nuclear counterstaining (blue). Dashed white lines delimit the dermal-epidermal junction and hair follicles. Images are representative of at least four mice per condition. Representative images are shown for each time point (n = 4 animals per genotype). Scale bar, 20 µm.

**Table S1.**
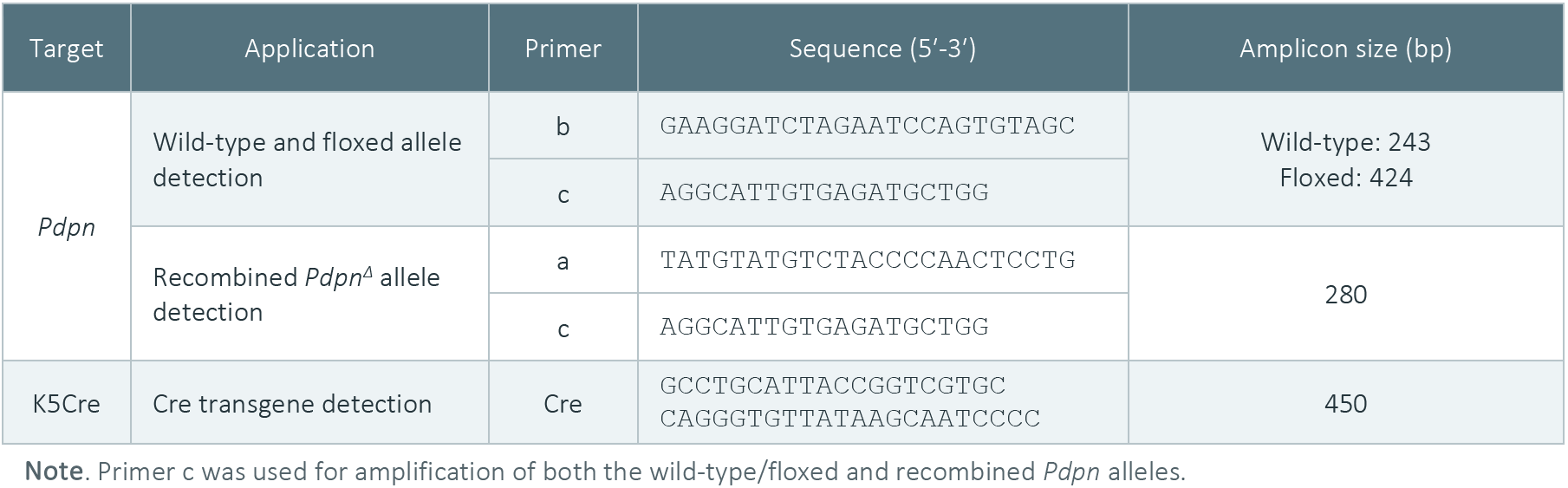
Oligonucleotides used for genotyping.

**Table S2.** Primary and secondary antibodies used in this study.

| Target | Clone | Host | Supplier | Catalogue no. | Dilution<br>(WB / IF) |
|---|---|---|---|---|---|
| **Primary antibodies** |  |  |  |  |  |
| Podoplanin | PA2.26 | Rat | Gandarillas et al., 1997 | — | 1:100 / 1:100 |
| Keratin 5 | Poly19055 | Rabbit | BioLegend | 905501 | 1:10,000 / 1:2,000 |
| Keratin 10 | Poly19054 | Rabbit | BioLegend | 905401 | 1:10,000 / 1:2,000 |
| Keratin 14 | Poly19053 | Rabbit | BioLegend | 905301 | — / 1:2,000 |
| Involucrin | Poly19244 | Rabbit | BioLegend | 924401 | 1:10,000 / 1:2,000 |
| Loricrin | Poly19051 | Rabbit | BioLegend | 905101 | 1:10,000 / 1:2,000 |
| Filaggrin | Poly19058 | Rabbit | BioLegend | 905801 | 1:10,000 / 1:2,000 |
| E-cadherin | ECCD-2 | Rat | Thermo Fisher Scientific | 13-1900 | 1:100 / 1:50 |
| Claudin-1 | JAY.8 | Rabbit | Thermo Fisher Scientific | 51-9000 | 1:100 / 1:500 |
| ZO-1 | — | Rabbit | Thermo Fisher Scientific | 40-2200 | 1:1,000 / 1:400 |
| BrdU | BU1/75 (ICR1) | Rat | Abcam | ab6326 | — / 1:250 |
| Ki67 | SP6 | Rabbit | Master Diagnostica (Vitro) | MAD-000310Q | — / 1:50 |
| α-Tubulin | DM1A | Mouse | Sigma-Aldrich | T6199 | 1:10,000 / — |
| β-Actin | AC-15 | Mouse | Sigma-Aldrich | A5441 | 1:10,000 / — |
| GAPDH | 6C5 | Mouse | Merck Millipore | MAB374 | 1:10,000 / — |
| β-Catenin | E247 | Rabbit | Abcam | ab32572 | — / 1:200 |
| β-Catenin | — | Mouse | BD Transduction Laboratories | 610154 | — / 1:200 |
| Desmoglein-1 | Clone 62 | Mouse | BD Transduction Laboratories | 610154 | 1:1,000 / — |
| Ezrin/Radixin/Moesin | — | Rabbit | Cell Signaling Technology | 3142 | 1:1,000 / - |
| Phospho-Ezrin (Thr567)/Radixin (Thr564)/Moesin (Thr558) | — | Rabbit | Cell Signaling Technology | 3141 | 1:1,000 / 1:100 |
| Phospho-Histone H3 (Ser10) | — | Rabbit | Merck Millipore | 06-570 | — / 1:250 |
| Myosin light chain 2 | — | Rabbit | Cell Signaling Technology | 3672 | 1:1,000 / - |
| Phospho-Myosin light chain 2 (Thr18/Ser19) | — | Rabbit | Cell Signaling Technology | 3674 | 1:1,000 / 1:100 |
| **Secondary antibodies** |  |  |  |  |  |
| Anti-mouse IgG–HRP | — | Sheep | Cytiva (Amersham) | NXA931V | 1:2,000 / — |
| Anti-rat IgG–HRP | — | Goat | Cytiva (Amersham) | NA935 | 1:5,000 / — |
| Anti-rabbit IgG–HRP | — | Donkey | Cytiva (Amersham) | NA934V | 1:5,000 / — |
| Anti-mouse IgG–Alexa Fluor 488 | — | Goat | Molecular Probes (Invitrogen) | A11029 | — / 1:200 |
| Anti-rat IgG–Alexa Fluor 488 | — | Goat | Molecular Probes (Invitrogen) | A11006 | — / 1:200 |
| Anti-rabbit IgG–Alexa Fluor 546 | — | Goat | Molecular Probes (Invitrogen) | A11035 | — / 1:200 |

*WB, western blotting; IF, immunofluorescence. (—), not applicable or not used, as appropriate. PA2.26 is a non-commercial antibody; their source is reported in the Supplier column. HRP, horseradish peroxidase*

## REFERENCES

1. Acton, S.E., Farrugia, A.J., Astarita, J.L., Mourão-Sá, D., Jenkins, R.P., Nye, E., Hooper, S., van Blijswijk, J., Rogers, N.C., Snelgrove, K.J., Rosewell, I., Moita, L.F., Stamp, G., Turley, S.J., Sahai, E., Reis e Sousa, C., 2014. Dendritic cells control fibroblastic reticular network tension and lymph node expansion. Nature 514, 498–502. 10.1038/nature13814

2. Asai, J., 2022. The Role of Podoplanin in Skin Diseases. Int. J. Mol. Sci. 23, 1310. 10.3390/ijms23031310

3. Astarita, J.L., Acton, S.E., Turley, S.J., 2012. Podoplanin: emerging functions in development, the immune system, and cancer. Front. Immunol. 3, 283. 10.3389/fimmu.2012.00283

4. Astarita, J.L., Cremasco, V., Fu, J., Darnell, M.C., Peck, J.R., Nieves-Bonilla, J.M., Song, K., Kondo, Y., Woodruff, M.C., Gogineni, A., Onder, L., Ludewig, B., Weimer, R.M., Carroll, M.C., Mooney, D.J., Xia, L., Turley, S.J., 2015. The CLEC-2–podoplanin axis controls the contractility of fibroblastic reticular cells and lymph node microarchitecture. Nat. Immunol. 16, 75–84. 10.1038/ni.3035

5. Baars, S., Bauer, C., Szabowski, S., Hartenstein, B., Angel, P., 2015. Epithelial deletion of podoplanin is dispensable for re-epithelialization of skin wounds. Exp. Dermatol. 24, 785–787. 10.1111/exd.12781

6. Box, K., Joyce, B.W., Devenport, D., 2019. Epithelial geometry regulates spindle orientation and progenitor fate during formation of the mammalian epidermis. eLife 8. 10.7554/eLife.47102

7. Cheok, Y.Y., Tan, G.M.Y., Chan, Y.T., Abdullah, S., Looi, C.Y., Wong, W.F., 2024. Podoplanin and its multifaceted roles in mammalian developmental program. Cells Dev. 203943. 10.1016/j.cdev.2024.203943

8. Cimini, M., Kishore, R., 2021. Role of Podoplanin-Positive Cells in Cardiac Fibrosis and Angiogenesis After Ischemia. Front. Physiol. 12, 667278. 10.3389/fphys.2021.667278

9. Cotsarelis, G., Sun, T.-T., Lavker, R.M., 1990. Label-retaining cells reside in the bulge area of pilosebaceous unit: Implications for follicular stem cells, hair cycle, and skin carcinogenesis. Cell 61, 1329–1337. 10.1016/0092-8674(90)90696-C

10. Damen, M., Wirtz, L., Soroka, E., Khatif, H., Kukat, C., Simons, B.D., Bazzi, H., 2021. High proliferation and delamination during skin epidermal stratification. Nat. Commun. 12, 3227. 10.1038/s41467-021-23386-4

11. Dlugosz, A.A., Yuspa, S.H., 1993. Coordinate changes in gene expression which mark the spinous to granular cell transition in epidermis are regulated by protein kinase C. J. Cell Biol. 120, 217–225. 10.1083/jcb.120.1.217

12. Durchdewald, M., Guinea-Viniegra, J., Haag, D., Riehl, A., Lichter, P., Hahn, M., Wagner, E.F., Angel, P., Hess, J., 2008. Podoplanin is a novel fos target gene in skin carcinogenesis. Cancer Res. 68, 6877–6883. 10.1158/0008-5472.CAN-08-0299

13. Gadre, P., Markova, P., Ebrahimkutty, M., Jiang, Y., Bouzada, F.M., Watt, F.M., 2024. Emergence and properties of adult mammalian epidermal stem cells. Dev. Biol. 515, 129–138. 10.1016/j.ydbio.2024.07.014

14. Gandarillas, A., Scholl, F.G., Benito, N., Gamallo, C., Quintanilla, M., 1997. Induction of PA2.26, a cell-surface antigen expressed by active fibroblasts, in mouse epidermal keratinocytes during carcinogenesis. Mol. Carcinog. 20, 10–18.

15. Gonzales, K.A.U., Fuchs, E., 2017. Skin and Its Regenerative Powers: An Alliance between Stem Cells and Their Niche. Dev. Cell 43, 387–401. 10.1016/j.devcel.2017.10.001

16. Hardman, M.J., Sisi, P., Banbury, D.N., Byrne, C., 1998. Patterned acquisition of skin barrier function during development. Development 125, 1541–1552.

17. Honma, M., Fujii, M., Iinuma, S., Minami-Hori, M., Takahashi, H., Ishida-Yamamoto, A., Iizuka, H., 2013. Podoplanin expression is inversely correlated with granular layer/filaggrin formation in psoriatic epidermis. J. Dermatol. 40, 296–297. 10.1111/1346-8138.12060

18. Honma, M., Minami-Hori, M., Takahashi, H., Iizuka, H., 2012. Podoplanin expression in wound and hyperproliferative psoriatic epidermis: regulation by TGF-β and STAT-3 activating cytokines, IFN-γ, IL-6, and IL-22. J. Dermatol. Sci. 65, 134–140. 10.1016/j.jdermsci.2011.11.011

19. Honma, M., Shibuya, T., Hayashi, K., Iinuma, S., Fujii, M., Ishida-Yamamoto, A., 2019. Suppression of podoplanin expression during differentiation of epidermal keratinocytes. J. Dermatol. 46, 922– 924. 10.1111/1346-8138.15017

20. Hur, Y.H., 2024. Epidermal stem cells: Interplay with the skin microenvironment during wound healing. Mol. Cells 47, 100138. 10.1016/j.mocell.2024.100138

21. Ishikawa, M., Phung, H.M., Dumrongphuttidecha, T., Sada, A., 2025. New insights into signaling networks coordinating epidermal stem cell regulation in skin regeneration and aging. Curr. Opin. Cell Biol. 97, 102594. 10.1016/j.ceb.2025.102594

22. Jiang, C., Javed, A., Kaiser, L., Nava, M.M., Xu, R., Brandt, D.T., Zhao, D., Mayer, B., Fernández-Baldovinos, J., Zhou, L., Höß, C., Sawmynaden, K., Oleksy, A., Matthews, D., Weinstein, L.S., Hahn, H., Gröne, H.-J., Graumann, P.L., Niessen, C.M., Offermanns, S., Wickström, S.A., Worzfeld, T., 2021. Mechanochemical control of epidermal stem cell divisions by B-plexins. Nat. Commun. 12, 1308. 10.1038/s41467-021-21513-9

23. Lechler, T., Fuchs, E., 2005. Asymmetric cell divisions promote stratification and differentiation of mammalian skin. Nature 437, 275–280. 10.1038/nature03922

24. Lowe, K.L., Finney, B.A., Deppermann, C., Hägerling, R., Gazit, S.L., Frampton, J., Buckley, C., Camerer, E., Nieswandt, B., Kiefer, F., Watson, S.P., 2015. Podoplanin and CLEC-2 drive cerebrovascular patterning and integrity during development. Blood 125, 3769–3777. 10.1182/blood-2014-09-603803

25. Luxenburg, C., Amalia Pasolli, H., Williams, S.E., Fuchs, E., 2011. Developmental roles for Srf, cortical cytoskeleton and cell shape in epidermal spindle orientation. Nat. Cell Biol. 13, 203–214. 10.1038/ncb2163

26. Luxenburg, C., Heller, E., Pasolli, H.A., Chai, S., Nikolova, M., Stokes, N., Fuchs, E., 2015. Wdr1-mediated cell shape dynamics and cortical tension are essential for epidermal planar cell polarity. Nat. Cell Biol. 17, 592–604. 10.1038/ncb3146

27. Mahtab, E.A.F., Wijffels, M.C.E.F., Van Den Akker, N.M.S., Hahurij, N.D., Lie-Venema, H., Wisse, L.J., Deruiter, M.C., Uhrin, P., Zaujec, J., Binder, B.R., Schalij, M.J., Poelmann, R.E., Gittenberger-De Groot, A.C., 2008. Cardiac malformations and myocardial abnormalities in podoplanin knockout mouse embryos: Correlation with abnormal epicardial development. Dev. Dyn. Off. Publ. Am. Assoc. Anat. 237, 847–857. 10.1002/dvdy.21463

28. Makris, S., Hari-Gupta, Y., Cantoral-Rebordinos, J.A., Martinez, V.G., Horsnell, H.L., Benjamin, A.C., Cinti, I., Jovancheva, M., Shewring, D., Nguyen, N., Winde, C.M. de, Denton, A., Acton, S.E., 2024. Lymph Node Fibroblast Phenotypes and Immune Crosstalk Regulated by Podoplanin Activity. 10.1101/2022.12.01.518753

29. Martín-Villar, E., Borda-d’Agua, B., Carrasco-Ramirez, P., Renart, J., Parsons, M., Quintanilla, M., Jones, G.E., 2015. Podoplanin mediates ECM degradation by squamous carcinoma cells through control of invadopodia stability. Oncogene 34, 4531–4544. 10.1038/onc.2014.388

30. Martín-Villar, E., Fernández-Muñoz, B., Parsons, M., Yurrita, M.M., Megías, D., Pérez-Gómez, E., Jones, G.E., Quintanilla, M., 2010. Podoplanin associates with CD44 to promote directional cell migration. Mol. Biol. Cell 21, 4387–4399. 10.1091/mbc.E10-06-0489

31. Martín-Villar, E., Megías, D., Castel, S., Yurrita, M.M., Vilaró, S., Quintanilla, M., 2006. Podoplanin binds ERM proteins to activate RhoA and promote epithelial-mesenchymal transition. J. Cell Sci. 119, 4541–4553. 10.1242/jcs.03218

32. Martín-Villar, E., Scholl, F.G., Gamallo, C., Yurrita, M.M., Muñoz-Guerra, M., Cruces, J., Quintanilla, M., 2005. Characterization of human PA2.26 antigen (T1alpha-2, podoplanin), a small membrane mucin induced in oral squamous cell carcinomas. Int. J. Cancer 113, 899–910. 10.1002/ijc.20656

33. Miroshnikova, Y.A., Le, H.Q., Schneider, D., Thalheim, T., Rübsam, M., Bremicker, N., Polleux, J., Kamprad, N., Tarantola, M., Wang, I., Balland, M., Niessen, C.M., Galle, J., Wickström, S.A., 2018. Adhesion forces and cortical tension couple cell proliferation and differentiation to drive epidermal stratification. Nat. Cell Biol. 20, 69–80. 10.1038/s41556-017-0005-z

34. Montero-Montero, L., Renart, J., Ramírez, A., Ramos, C., Shamhood, M., Jarcovsky, R., Quintanilla, M., Martín-Villar, E., 2020. Interplay between Podoplanin, CD44s and CD44v in Squamous Carcinoma Cells. Cells 9, 2200. 10.3390/cells9102200

35. Quintanilla, M., Montero-Montero, L., Renart, J., Martín-Villar, E., 2019. Podoplanin in Inflammation and Cancer. Int. J. Mol. Sci. 20. 10.3390/ijms20030707

36. Ramirez, A., Page, A., Gandarillas, A., Zanet, J., Pibre, S., Vidal, M., Tusell, L., Genesca, A., Whitaker, D.A., Melton, D.W., Jorcano, J.L., 2004. A keratin K5Cre transgenic line appropriate for tissue-specific or generalized Cre-mediated recombination. Genes. N. Y. N 2000 39, 52–57. 10.1002/gene.20025

37. Rübsam, M., Mertz, A.F., Kubo, A., Marg, S., Jüngst, C., Goranci-Buzhala, G., Schauss, A.C., Horsley, V., Dufresne, E.R., Moser, M., Ziegler, W., Amagai, M., Wickström, S.A., Niessen, C.M., 2017. E-cadherin integrates mechanotransduction and EGFR signaling to control junctional tissue polarization and tight junction positioning. Nat. Commun. 8, 1250. 10.1038/s41467-017-01170-7

38. Schacht, V., Ramirez, M.I., Hong, Y.-K., Hirakawa, S., Feng, D., Harvey, N., Williams, M., Dvorak, A.M., Dvorak, H.F., Oliver, G., Detmar, M., 2003. T1alpha/podoplanin deficiency disrupts normal lymphatic vasculature formation and causes lymphedema. EMBO J. 22, 3546–3556. 10.1093/emboj/cdg342

39. Schmitz, A., Lazi’ć, E., Koumaki, D., Kuonen, F., Verykiou, S., Rübsam, M., 2015. Assessing the In Vivo Epidermal Barrier in Mice: Dye Penetration Assays. J. Invest. Dermatol. 135, 1–4. 10.1038/jid.2014.495

40. Scholl, F.G., Gamallo, C., Vilaró, S., Quintanilla, M., 1999. Identification of PA2.26 antigen as a novel cell-surface mucin-type glycoprotein that induces plasma membrane extensions and increased motility in keratinocytes. J. Cell Sci. 112 ( Pt 24), 4601–4613.

41. Sesartić, M., Ikenberg, K., Yoon, S.-Y., Detmar, M., 2020. Keratinocyte-Expressed Podoplanin is Dispensable for Multi-Step Skin Carcinogenesis. Cells 9, 1542. 10.3390/cells9061542

42. Shibuya, T., Honma, M., Fujii, M., Iinuma, S., Ishida-Yamamoto, A., 2019. Podoplanin suppresses the cell adhesion of epidermal keratinocytes via functional regulation of β1-integrin. Arch. Dermatol. Res. 311, 45–53. 10.1007/s00403-018-1878-9

43. Simpson, C.L., Patel, D.M., Green, K.J., 2011. Deconstructing the skin: cytoarchitectural determinants of epidermal morphogenesis. Nat. Rev. Mol. Cell Biol. 12, 565–580. 10.1038/nrm3175

44. Sotiropoulou, P.A., Candi, A., Blanpain, C., 2008. The Majority of Multipotent Epidermal Stem Cells Do Not Protect Their Genome by Asymmetrical Chromosome Segregation. Stem Cells 26, 2964– 2973. 10.1634/stemcells.2008-0634

45. Sumigray, K.D., Lechler, T., 2015. Cell adhesion in epidermal development and barrier formation. Curr. Top. Dev. Biol. 112, 383–414. 10.1016/bs.ctdb.2014.11.027

46. Sun, X., Joost, S., Kasper, M., 2023. Plasticity of Epithelial Cells during Skin Wound Healing. Cold Spring Harb. Perspect. Biol. 15, a041232. 10.1101/cshperspect.a041232

47. The landscape of ‘podoplanin’ in dermatology : Journal of the Egyptian Women’s Dermatologic Society [WWW Document], n.d. . Ovid. 10.4103/jewd.jewd_18_21

48. Toll, A., Gimeno-Beltrán, J., Ferrandiz-Pulido, C., Masferrer, E., Yébenes, M., Jucglà, A., Abal, L., Martí, R.M., Sanmartín, O., Baró, T., Casado, B., Gandarillas, A., Barranco, C., Costa, I., Mojal, S., García-Patos, V., Pujol, R.M., 2012. D2-40 immunohistochemical overexpression in cutaneous squamous cell carcinomas: A marker of metastatic risk. J. Am. Acad. Dermatol. 67, 1310–1318. 10.1016/j.jaad.2012.03.007

49. Tunggal, J.A., Helfrich, I., Schmitz, A., Schwarz, H., Günzel, D., Fromm, M., Kemler, R., Krieg, T., Niessen, C.M., 2005. E-cadherin is essential for in vivo epidermal barrier function by regulating tight junctions. EMBO J. 24, 1146–1156. 10.1038/sj.emboj.7600605

50. Verdoni, A.M., Ikeda, S., Ikeda, A., 2010. Serum response factor is essential for the proper development of skin epithelium. Mamm. Genome 21, 64–76. 10.1007/s00335-009-9245-y

51. Williams, S.E., Beronja, S., Pasolli, H.A., Fuchs, E., 2011. Asymmetric Cell Divisions Promote Notch-Dependent Epidermal Differentiation. Nature 470, 353–358. 10.1038/nature09793

52. Yin, H., Hu, M., Li, D., 2023. Regulation of epidermal stratification and development by basal keratinocytes. J. Cell. Physiol. 238, 742–748. 10.1002/jcp.30978

53. Yoon, S.-Y., Dieterich, L.C., Tacconi, C., Sesartic, M., He, Y., Brunner, L., Kwon, O., Detmar, M., 2019. An important role of podoplanin in hair follicle growth. PLOS ONE 14, e0219938. 10.1371/journal.pone.0219938

